# XENTURION, a multidimensional resource of xenografts and tumoroids from metastatic colorectal cancer patients for population-level translational oncology

**DOI:** 10.1101/2023.07.10.548375

**Authors:** Simonetta M. Leto, Elena Grassi, Marco Avolio, Valentina Vurchio, Francesca Cottino, Martina Ferri, Eugenia R. Zanella, Laura di Blasio, Desiana Somale, Marianela Vara-Messler, Francesco Galimi, Francesco Sassi, Barbara Lupo, Irene Catalano, Marika Pinnelli, Marco Viviani, Luca Sperti, Alfredo Mellano, Alessandro Ferrero, Caterina C. Zingaretti, Alberto Puliafito, Luca Primo, Andrea Bertotti, Livio Trusolino

## Abstract

The breadth and depth at which cancer models are interrogated contribute to successful translation of drug discovery efforts to the clinic. In colorectal cancer (CRC), model availability is limited by a dearth of large-scale collections of patient-derived xenografts (PDXs) and paired tumoroids from metastatic disease, the setting where experimental therapies are typically tested. XENTURION is a unique open-science resource that combines a platform of 129 PDX models and a sister platform of 129 matched PDX-derived tumoroids (PDXTs) from patients with metastatic CRC, with accompanying multidimensional molecular and therapeutic characterization. A PDXT-based population trial with the anti-EGFR antibody cetuximab revealed variable sensitivities that were consistent with clinical response biomarkers, mirrored tumor growth changes in matched PDXs, and recapitulated the outcome of EGFR genetic deletion. Adaptive signals upregulated by EGFR blockade were computationally and functionally prioritized, and inhibition of top candidates increased the magnitude of response to cetuximab. These findings illustrate the probative value and accuracy of large *ex vivo* and *in vivo* living biobanks, highlight the importance of cross-platform and cross-methodology systematic validation, and offer avenues for molecularly informed preclinical research.

## INTRODUCTION

Historically, the development of cancer organoids (tumoroids) from CRC samples has paved the way for the establishment of living biobanks of different tumor types^1^. Several studies have documented the practicability of deploying CRC tumoroids for drug screens using approved or investigational compounds, with drug sensitivities often dictated by underlying genetic susceptibilities^2–10^. For example, pharmacologic assays in tumoroids revealed an association between loss of function of the WNT pathway negative regulator RNF43 and exquisite sensitivity to chemical blockade of the O-acyltransferase porcupine, which is required for proper secretion of WNT morphogens^1^. CRC tumoroids proved valuable also for the identification of therapeutic opportunities with off-label compounds, such as repurposing of the antimitotic agent paclitaxel to CRC tumors with CpG island hypermethylation and epigenetic silencing of the E3 ubiquitin-protein ligase CHFR^9^. In a translational perspective, the potential and feasibility of using tumoroids as ‘avatars’ to guide individualized decision-making have been supported by proof-of-concept co-clinical trials, which showed a relatively high concordance between *ex vivo* drug sensitivities measured in CRC tumoroids and responses to the same treatments experienced by donor patients^5,11–13^.

Although the initial implementation of tumoroid technology has contributed to improving precision medicine in CRC^14–16^, the long-term impact of CRC tumoroid collections has lagged expectations in terms of numbers of models generated, extent and depth of molecular profiling, clinical representativeness, and value in predicting therapeutic response at the population level. The CRC tumoroid platforms established so far typically include less than a hundred samples. The composition of existing catalogs is further reduced to the order of tens when considering the availability of accompanying molecular and pharmacologic information, and drops down to only a handful of models that also have matched PDX counterparts for running companion *in vivo* studies^17,18^. Owing to this paucity of cases, current biobanks of CRC tumoroids have important limitations: i) they often fail to reflect intertumor diversity, which weakens the predictive power of regression models when trying to extract subgroup-defined genotype/phenotype associations; ii) they hardly contemplate systematic *in vivo* validation with paired xenografts, which diminishes the translational value of tumoroid-based drug development pipelines; iii) small sample size also complicates methodological work, such as assessing whether tumoroid establishment is biased by biological traits in originating tumors that influence cell culture viability.

Another aspect that challenges the clinical transferability of drug screen approaches using CRC tumoroids is that, with some exceptions^5–7,11^, pharmacologic experiments have been typically conducted in cultures established from colon primary tumor samples from treatment-naïve patients. This may cause attrition in drug development, as investigational compounds that enter the clinical space after successful preclinical testing are almost invariably administered to heavily pretreated metastatic patients. Finally, there is growing appreciation of the importance of standardizing practices and protocols to enhance reproducibility^19^. This need calls for ordered procedural attempts to optimize culture conditions, validate source characteristics, and define the most reliable measures of effects and the most accurate endpoint methodologies.

To tackle some of these hurdles we have developed XENTURION, a resource of matched XENografts and TUmoroids for Research In ONcology that encompasses 129 sibling pairs of PDXs and PDXTs from patients with metastatic CRC. The vast majority of XENTURION models underwent comparative analysis at the mutational, gene copy number and transcriptomic levels and were annotated for response to the clinically approved anti-EGFR antibody cetuximab *ex vivo* and *in vivo*, showing substantial consistency in all the data levels examined. Further, we embarked on a proof-of-principle discovery effort to identify and prioritize druggable co-extinction targets, and found that inhibition of top candidates increased the depth of response to cetuximab. This platform addresses a long-standing quest for large-scale collections of extensively annotated CRC preclinical models for integrative *ex vivo* and *in vivo* translational applications. All molecular profiles and therapeutic annotations are accessible in public repositories and as Supplementary Data here, and models are available for distribution to non-profit entities. By accessing XENTURION, the biomedical community will have at their disposal a knowledge base of disseminatable methods, resources and information to streamline preclinical studies and accelerate new treatments for patients with advanced CRC.

## RESULTS

### Facts and figures of XENTURION

The workflow of XENTURION characterization is illustrated in Fig. 1A. We established a biobank of matched PDXs and PDXTs, representative of the biological and clinical diversity of metastatic CRC, in which PDXTs were quality-checked at multiple levels to ascertain whether they can be used as faithful proxies of parental PDXs to filter, skim and prioritize anticancer agents in the development pipeline prior to more laborious *in vivo* validation. Most of existing CRC tumoroid collections have been generated directly from original tumors in patients, and little is known about the molecular and biological fidelity of PDXTs. To fill this gap, we performed a systematic comparison between paired PDXs and PDXTs in terms of mutational profiles, gene copy number architecture, transcriptomic features, and responsiveness to standard-of-care therapy. This comparative effort was also leveraged to extract genes that were concordantly modulated by drug pressure in PDXs and PDXTs, with the aim to identify hits potentially involved in tumor adaptation to therapeutic stress. Following hit nomination, a stepwise drug screen for actionable targets was conducted in PDXTs, and surviving candidate compounds were finally tested *in vivo*.

**Figure 1.**
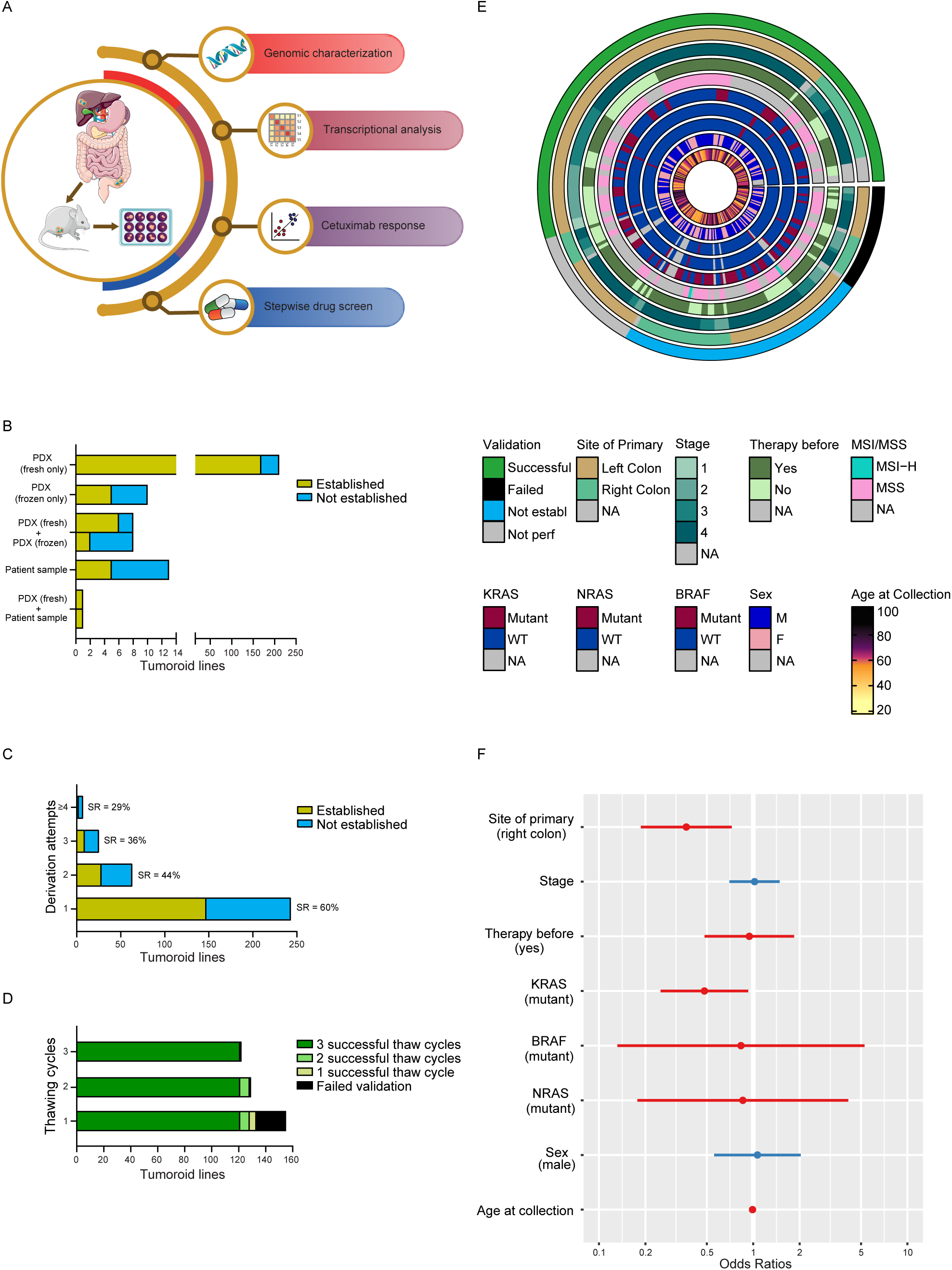
Facts and Figures of XENTURION. **A,** Schematic overview of XENTURION experimental design. Matched PDXTs and PDXs were subjected to comparative mutational, gene copy number and transcriptomic analyses. Molecular annotation was paralleled by systematic assessment of *ex vivo* and *in vivo* response to cetuximab. Post-cetuximab transcriptomic profiles were leveraged to extract upregulated genes potentially involved in adaptive resistance to EGFR blockade. Compounds against candidate targets were tested in a stepwise drug screen, and those that proved effective in PDXT assays underwent final validation in PDXs. **B, C,** Success rate in the establishment of tumoroid lines according to the nature of the sample of origin (**B**) or the number of derivation attempts (**C**). When tumoroids were established from different originating samples (e.g., fresh and frozen PDX explants), success rates were computed for models derived from freshly explanted tumors. SR, success rate. **D,** Number of validated tumoroids according to the number of freeze-thaw cycles. **E,** Main clinical and molecular features of the starting population from which tumoroid derivation was attempted. The circus plot includes all cases with successful validation (*N* = 133), those that failed validation (*N* = 24), established cases for which validation was not performed (Not perf, *N* = 29), and those that failed initial establishment (Not establ, *N* = 57). F, female; M, male; MSI-H, microsatellite instability high; MSS, microsatellite stability; NA, not available; WT, wild-type. **F**, Odds ratios of a multivariate logistic regression with PDXT establishment/validation success status (1, successful, *N* = 129; 0, failed, *N* = 73) as dependent variable and several clinical and molecular annotations as independent variables. Red color indicates that the independent variable has a negative effect on the validation rate; blue color indicates the opposite. The only continuous variables are stage and age at collection; all other variables are binary.

Between March 2015 and September 2021, a total of 267 CRC liver metastases from 264 patients were processed for tumoroid derivation (for three cases, two liver metastases from the same patient were available). The clinical history of donor patients is summarized in Supplementary Table 1. To minimize alterations in the biology of tumors and avoid biased selection of specific growth dependencies, we standardized culture conditions that sustained long-term growth of tumoroids in a medium with minimal composition, in line with the notion that CRC tumoroids become gradually independent from niche signals during cancer progression^20^. For preliminary inclusion into the biobank, each PDX-tumoroid pair had to show matching identity by genetic fingerprinting, negativity for human and mouse pathogens, and a histology congruent with CRC phenotypes. This approach left us with 243 models; 19 cases were excluded because of wrong fingerprinting; four were diagnosed as lymphomas by histopathological evaluation; and one was excluded for technical reasons (deterioration of archived material) (Supplementary Table 2).

We defined tumoroids as ‘established’ when they could be propagated for at least three passages or expanded enough to be cryopreserved. The vast majority of samples (211/243, 87%) were processed using freshly explanted PDX tumors as only source, with an 80% establishment success rate (169/211) (Fig. 1B and Supplementary Table 2). In the few cases where PDXT derivation was attempted starting from frozen PDX tumors, the success rate was lower (5/10, 50%) (Fig. 1B and Supplementary Table 2). Differences in PDXT establishment were also observed when tumoroids were derived in parallel from fresh and frozen material from the same PDX; in particular, starting from eight PDXs for which both fresh and frozen tumor fragments were available, the establishment rate was 75% for fresh tissues (6/8), and 25% for frozen material (2/8) (Fig. 1B and Supplementary Table 2). Although the number of PDXTs established from frozen PDX tumors is small, these figures suggest that freshly explanted tumors may be more suitable to PDXT establishment than frozen material. XENTURION also includes 13 tumoroids derived directly from fresh human specimens after surgery; in this subgroup, the success rate in the production of established tumoroids was markedly lower (5/13, 38%). Finally, for one case, tumoroids were successfully established from both fresh PDX explants and the original patient sample (Fig. 1B and Supplementary Table 2). Overall, XENTURION comprises 186 established tumoroids of metastatic CRC, with a success rate of 77% (186/243); the collection is almost completely represented by PDXT lines (181/186, 97%) for which paired PDXs are available (Fig. 1B and Supplementary Table 2). For tumoroid establishment, a single derivation procedure was sufficient in most models, with a success rate of 60% (147/243) (Fig. 1C). When establishment failed after the first attempt, two or more additional rounds were performed if PDXs were in line. The success rate of tumoroid establishment decreased proportionally with attempt repetition, specifically to 44% after the second attempt, 36% after the third attempt, and 29% after the fourth or subsequent attempts (Fig. 1C). Hence, we can reasonably conclude that tumoroids of metastatic CRC that do not grow in culture after the first derivation round are less likely to become established models.

Established tumoroids were credentialed as bona fide immortalized models, capable of long-term recovery and expansion, through a stringent validation protocol that included periodic identity checks, microbiologic tests and various freeze-thaw cycles. One hundred and forty-five established tumoroids underwent at least three freeze-thaw cycles, and 121 (83%) passed validation (Fig. 1D and Supplementary Table 2). Of practical utility, lack of recovery after the first freeze-thaw cycle was sufficient to identify 92% (22/24) of cases that would not survive additional ‘rescue’ cycles, thus failing validation; at the same time, 100% of tumoroids that were recovered after the first freeze-thaw cycle successfully completed validation in subsequent cycles (Fig. 1D and Supplementary Table 2). For this reason, we relaxed the validation criteria and admitted in the final collection additional models that, at the time of manuscript writing, had survived two freeze-thaw cycles (seven cases) or one cycle (five cases). Ultimately, XENTURION encompasses a total of 133 validated tumoroids: 129 PDXTs (with paired PDXs) and four tumoroids directly derived from donor patients (Fig. 1B-D and Supplementary Table 2).

Fig. 1E summarizes the main clinical and molecular features of the samples that fed into XENTURION, including primary tumor sidedness and stage, patients’ sex, age and exposure to therapy before sample donation, DNA microsatellite status, and the presence of clinically relevant driver mutations. To explore whether tumoroid derivation favored over- or under-representation of such features in XENTURION with respect to the starting population, we assessed their relative distribution in validated models versus those that failed establishment or validation. Enrichment analysis revealed that the establishment or validation of metastatic samples with primary tumor location in the right colon was unsuccessful more often than expected by chance (Fig. 1F). This could be due to the fact that left-sided CRC tumors are usually more dependent on EGFR signaling^21^, thus more stimulated to grow by the EGF ligand present in the culture medium, than right-sided tumors. A similar enrichment among samples that failed validation was observed for tumors harboring *KRAS* mutations (Fig. 1F). This was quite unexpected, as *KRAS* mutant CRC tumors are notoriously more aggressive than *KRAS* wild-type tumors^22,23^ and ectopic introduction of mutant *KRAS* promotes – rather than contrasts – the expansion of CRC tumoroids^24^. We suspected that the higher representation of *KRAS* mutant cases among tumoroids that did not pass validation could be due to a procedural bias related to the time when tumoroids were generated. PDXTs from *KRAS* wild-type tumors were more often derived from late-passage (more than three) PDXs, typically from large cohorts that had been propagated *in vivo* several times to obtain an adequate number of replicas for testing with the anti-EGFR antibody cetuximab. Since mutant *KRAS* is known to confer resistance to cetuximab^25^, PDXs with *KRAS* mutations were not repeatedly expanded for cetuximab treatment, and tumoroids were generated from smaller cohorts at earlier passages. Confirming the hypothesis that PDX passaging rather than *KRAS* mutations impacted on PDXT stability, we found that late-passage PDXs were more likely to give rise to validated tumoroids than early-passage PDXs (Supplementary Fig. 1). This is in line with our observation that fresh samples from patients (never passaged in mice) were less prone to grow in culture (Fig. 1B) and suggests that serial mouse engraftment eases adaptation of cancer cells to long-term propagation *ex vivo*.

### Mutational and gene copy number analysis of paired PDXTs and PDXs reveals substantial model concordance

We performed targeted next-generation sequencing of 116 relevant CRC genes^26^ to detect small somatic alterations [single nucleotide variants (SNVs) and indels] in a set of 144 PDXTs and their matched PDXs. The overall distribution of allele frequencies and the number of identified variants were consistent between PDXTs and PDXs (Supplementary Fig. 2). Mutational profiles were analyzed more in depth for a subset of 125 sibling pairs, in which only validated PDXTs were included. At the level of individual genes, the vast majority of mutations were conserved, with no preferential occurrence in PDXTs or PDXs (Fig. 2A). This consistency was maintained also at the level of specific mutations; indeed, the Jaccard similarity coefficient was markedly higher for matched models than for unmatched models (average matched, 0.828; average unmatched, 0.006) (Fig. 2B). Importantly, the extent of mutational concordance between PDXTs and PDXs was superimposable to that of a recent comparison of 536 original patient tumors and matched PDXs across 25 cancer types^27^, indicating negligible divergence between pre-derivation samples, PDXs and PDXTs when considering the general mutational repertoire.

**Figure 2.**
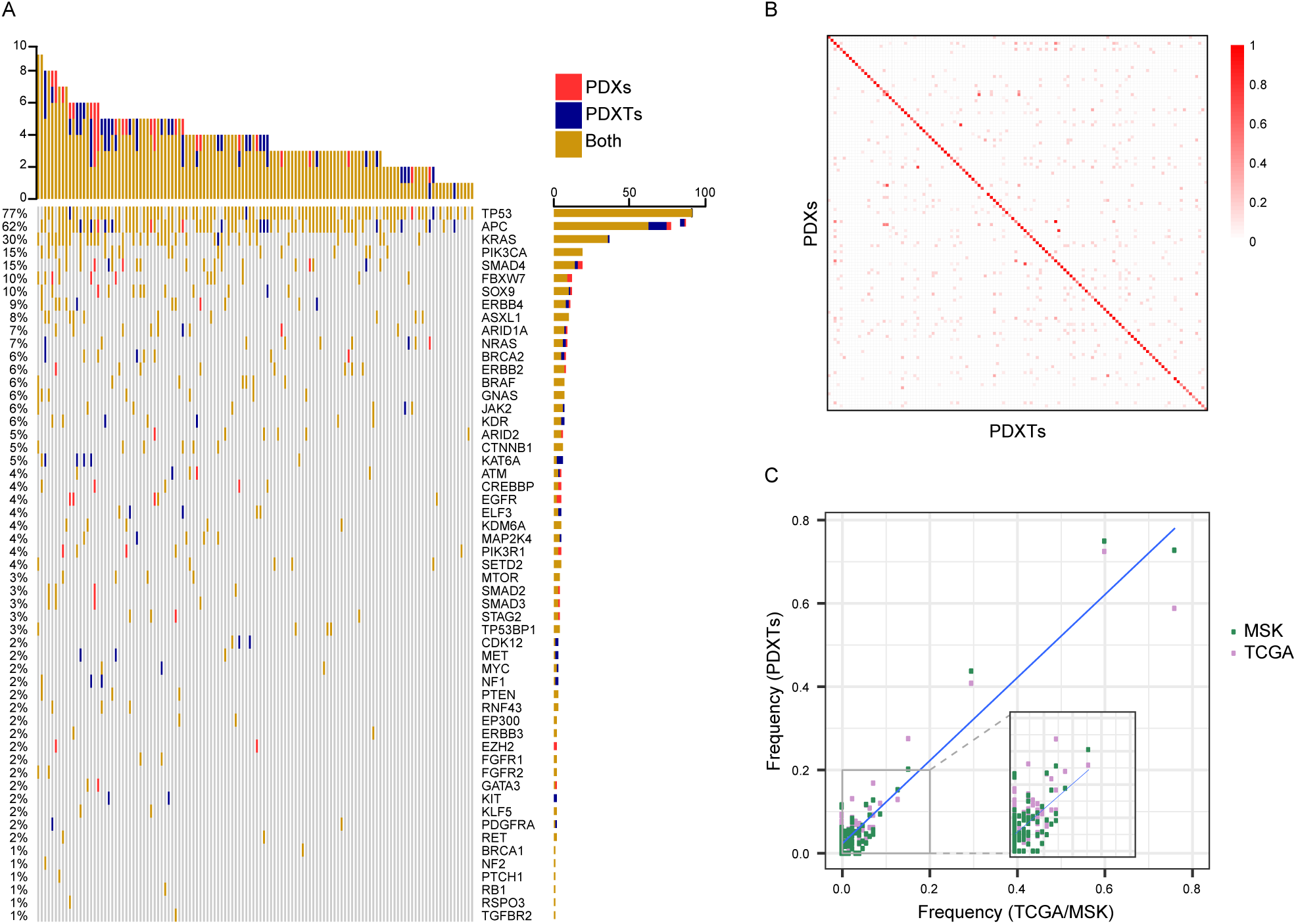
Comparative landscape of somatic single nucleotide variations and indels in paired PDXTs and PDXs. **A,** Common and private alterations in 124 pairs of matched PDXTs and PDXs. One pair for which mutational data were available was excluded because no alterations with allele frequencies > 0.05 were detected. Genes without any alteration in the whole cohort were removed. The top barchart shows the total number of mutations for each sample. The barchart on the right shows the percentage of mutations for each gene in the cohort. **B,** Jaccard similarity indexes of somatic alterations between 124 matched PDXs and PDXTs. *P* < 2.2e-308 by two tailed Mann-Whitney test. **C,** Gene-level population frequencies of mutational alterations in PDXTs versus those detected in the TCGA dataset or the MSK-IMPACT dataset; the inset shows that the correlation is not driven solely by genes with high mutational frequencies. Pearson coefficient, 0.93 (*P* = 1.46e-51) for TCGA; Pearson coefficient, 0.96 (*P* = 2.37e-64) for MSK-IMPACT.

Subclonal variants that are poorly represented in PDXs may become dominant in PDXTs if they confer a growth advantage in culture. To study the clonal composition of PDX-PDXT pairs, we evaluated the representation of shared mutations with allele frequencies higher than 0.05 for genes with more than five alterations in the collection. The 0.05 threshold was deliberately set low to gauge the impact of tumoroid derivation on tumor subclonal architecture. No significant differences in allele frequencies emerged from this analysis (Supplementary Fig. 3), indicating that intratumor clonal heterogeneity was substantially preserved in PDXTs with respect to originating PDXs. In some cases, the allele frequencies of alterations in frequently mutated tumor suppressor genes (for example, *APC* and *TP53*) were 1 in PDXTs and slightly lower in the paired xenografts (Supplementary Fig. 3), suggesting subtle defects in the filtering procedure of mouse reads deriving from host stromal contamination.

Next, we compared the frequency of gene alterations in our collection with that of two large datasets of human samples from CRC patients: TCGA, which mainly includes primary tumors^26^, and MSK-IMPACT, which is predominantly composed of metastatic samples^28^. We found significant correlations between PDXTs and the two clinical datasets (Pearson coefficient, 0.93 for TCGA and 0.96 for MSK-IMPACT) (Fig. 2C) as well as between PDXs and the clinical datasets (Pearson coefficient, 0.92 for TCGA and 0.95 for MSK-IMPACT) (Supplementary Fig. 4). This comparison indicates that XENTURION reflects the mutational landscape of patient cohorts and points to a substantial similarity of mutational frequencies in primary and metastatic CRC tumors.

We then investigated whether the PDXT validation protocol may result in the enrichment or depletion of defined variants. We applied univariate logistic regression models to predict validation using the entire XENTURION sequencing dataset, which includes mutational data also for 7 non-validated PDXTs. We considered the presence or absence of any SNVs or indels in PDXTs as independent variables, and the validation status as dependent variable. When testing genes mutated in at least five tumoroids, only mutations in the *CTNNB1* gene (encoding β-catenin) were significantly over­represented in PDXTs that failed validation (odds ratio, 0.067) (Supplementary Fig. 5 and Supplementary Table 3). Both *CTNNB1* and *APC* mutations result in constitutive activation of the Wnt pathway (which sustains CRC proliferation), but mutant β-catenin is known to be more modulatable by exogenous Wnt stimulation than mutant APC^29^. Since PDXTs were cultured in the absence of Wnt agonists, it is conceivable that *CTNNB1* mutant samples are less fit to grow in a nutrient-poor medium than *APC* mutant samples.

Most colorectal tumors display chromosomal instability, which could be exacerbated by evolutionary bottlenecks such as those imposed by tissue culture propagation. To examine whether copy number changes materialized in our models following *ex vivo* culturing, we surveyed PDXTs versus their matched PDXs using DNA shallow sequencing in the same 125 pairs used for mutational profiling. We found a high consistency of copy number variations between PDXTs and the corresponding PDXs compared with unmatched samples, as shown by Pearson correlations between the segmented log ratios (average matched, 0.88; average unmatched, 0.39) (Fig. 3A). In PDXTs, the overall landscape of chromosomal alterations was in line with that observed in patients^26^. In particular, whole-arm copy number gains were detected in chromosomes 7, 13 and 20, and long-arm specific gains were detected in chromosome 1 and 8; whole-arm losses occurred in chromosome 18 (where the *SMAD4* gene lies) and in the short arms of chromosomes 1 and 8 (Fig. 3B). Accordingly, the population frequencies of copy number alterations at the gene level, obtained with GISTIC, were positively correlated between PDXTs and the TCGA or MSK-IMPACT patient cohorts [Pearson coefficient, 0.92 (gains) and 0.87 (losses) for TCGA; 0.80 (gains) and 0.88 (losses) for MSK-IMPACT] (Fig. 3C) and between PDXs and the patient cohorts [Pearson coefficient, 0.92 (gains) and 0.90 (losses) for TCGA; 0.84 (gains) and 0.89 (losses) for MSK-IMPACT] (Supplementary Fig. 6). In summary, our data suggest that PDXTs generally retain the mutational and genomic structure of parental PDXs. Moreover, the distribution of major mutational drivers and copy number alterations observed in XENTURION PDXTs and PDXs is largely superimposable to that of human CRC samples.

**Figure 3.**
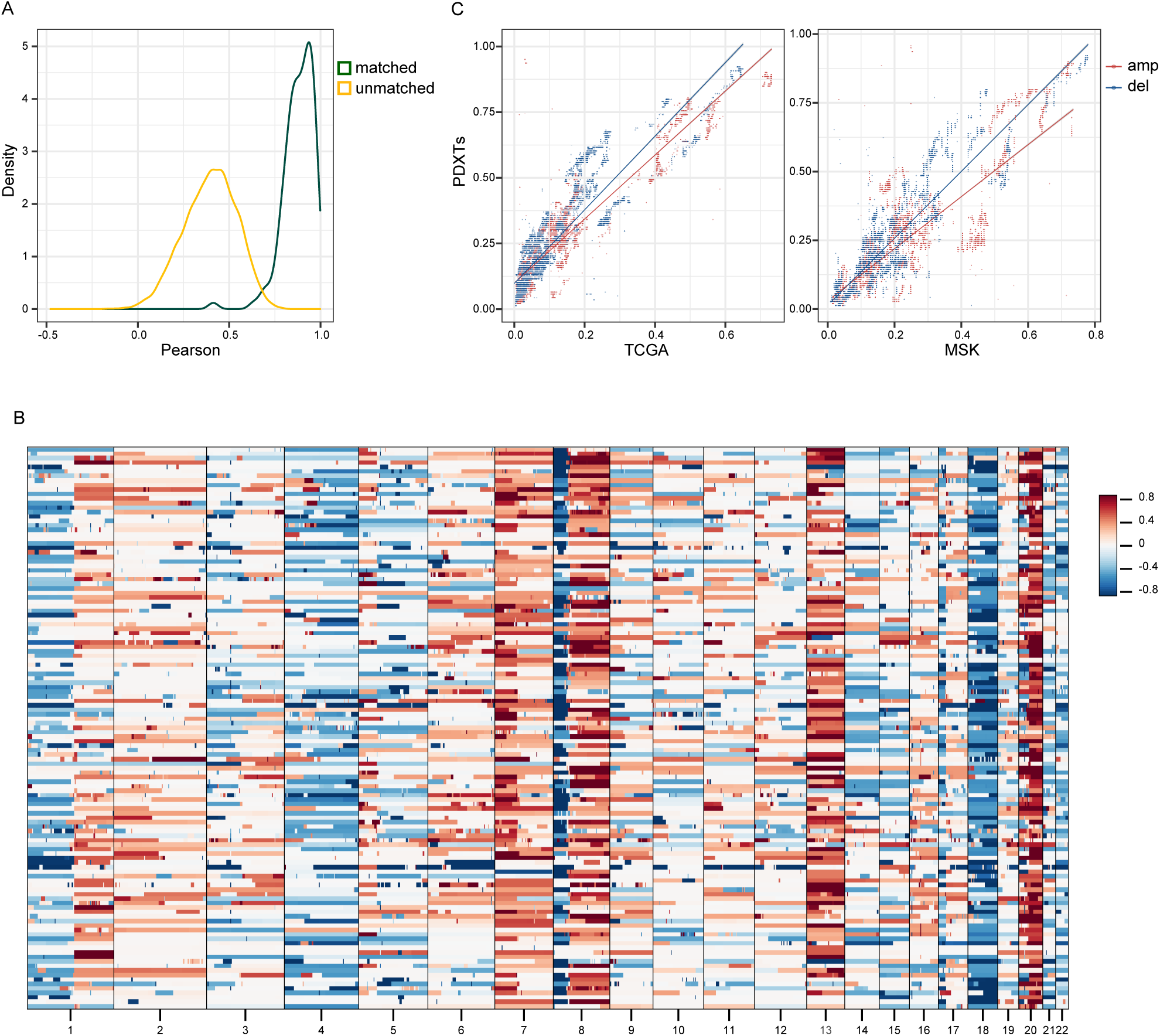
Comparative copy number architecture in paired PDXTs and PDXs. **A,** Distribution of Pearson correlations between copy number profiles of matched (*N* = 125) and unmatched (*N* = 15,500) pairs of PDXTs and PDXs. Matched pairs, average Pearson coefficient, 0.88; unmatched pairs, average Pearson coefficient 0.39; matched versus unmatched pairs, *P* = 1.92e-81 by two-tailed Mann-Whitney test. **B,** Autosomal copy number profiles of PDXTs (*N* = 125), expressed in segmented log2 ratio of the normalized read depth. Red and blue colors indicate gain and loss events, respectively. **C,** Gene-level population frequencies of gain or loss events, as identified by GISTIC, in PDXTs versus those detected in the TCGA or the MSK-IMPACT datasets. Pearson coefficient, 0.92 (gains) and 0.87 (losses) for TCGA; 0.80 (gains) and 0.88 (losses) for MSK-IMPACT (*P* < 1e-230 for both comparisons). Amp, amplification; del, deletion.

### PDXT transcriptional identity retains fidelity to corresponding PDXs

RNAseq data were used to compare the transcriptomic phenotypes of 21 surgical specimens from donor patients (human liver metastases, HLMs), 119 PDXs, and 124 PDXTs. Based on gene ontology (GO) enrichment analysis of differentially expressed genes, gene signatures related to cellular division and DNA replication were more abundant in PDXs than HLMs, consistent with the faster growth rates of xenografts compared with those of tumors in patients^30^ (Supplementary Table 4). Pathways downregulated in PDXs versus HLMs were associated with innate and adaptive immunity and stromal remodeling (Supplementary Table 4), as expected for models grown in immunocompromised animals and in agreement with the observation that human stromal cells are replaced by mouse components soon after tumor implantation^31,32^. Being derived from PDXs, PDXTs predictably showed similar upregulated and downregulated pathways with respect to HLMs (Supplementary Table 4).

Comparative analysis of xenografts and tumoroids revealed that gene signatures of steroid, retinoid and fatty acid metabolism were more expressed in PDXTs than in PDXs (Supplementary Fig. 7 and Supplementary Table 4). This might be due to metabolic adaptations to the culture conditions and is in line with previous results obtained in a smaller set of 19 CRC PDX/tumoroid sibling pairs^4^. A cluster of gene sets functionally related to ribosome biogenesis and protein translation stood out as significantly downregulated in PDXTs versus PDXs (Supplementary Fig. 7 and Supplementary Table 4). We surmise that these traits of protein synthesis impediment are ascribable to the hyperoxic environment experienced by tumoroids in standard culture conditions, as it is known that hyperoxic conditions suppress mRNA translation^33,34^.

To investigate transcriptional fidelity of PDXT-PDX pairs, we first considered a subset of 79 ‘super­matched’ samples selected based on genealogical proximity (*i.e.*, pairs made of an early-passage PDXT with its nearest ancestor PDX). This analysis showed high consistency in transcript abundance between samples, with an intramodel Pearson correlation significantly greater than intermodel correlations (average matched, 0.82; average unmatched, 0.61) (Fig. 4A). We then extended the survey to a more variegate set of sample families, which included early and late propagations from the same PDXT and one or more matched PDXs grown in more distant generations of mice. In this larger set, consisting of 116 models for a total of 308 biological replicates, the similarity between matched PDXTs and PDXs was confirmed; in particular, the intramodel Pearson correlation was significantly higher than intermodel correlations (average matched, 0.78; average unmatched, 0.52), and 80/116 (69%) models derived from the same originating tumor proved to belong to the same cluster by unsupervised hierarchical clustering (Supplementary Fig. 8). We note that the transcriptional profile of tumor CRC1241, which had a very high PDXT-PDX correlation (Pearson coefficient, 0.90), was different from the rest of our cohort (average Pearson coefficient, 0.31) (Fig. 4A). Accordingly, a deep-learning tool that uses RNA gene expression data to infer a tumor’s primary tissue of origin^35^ predicted the tumor as a cervical squamous cell carcinoma (Supplementary Table 5), and retrospective pathological revision cataloged it as an anal squamous cell carcinoma. For this reason, we excluded CRC1241 from further analyses.

**Figure 4.**
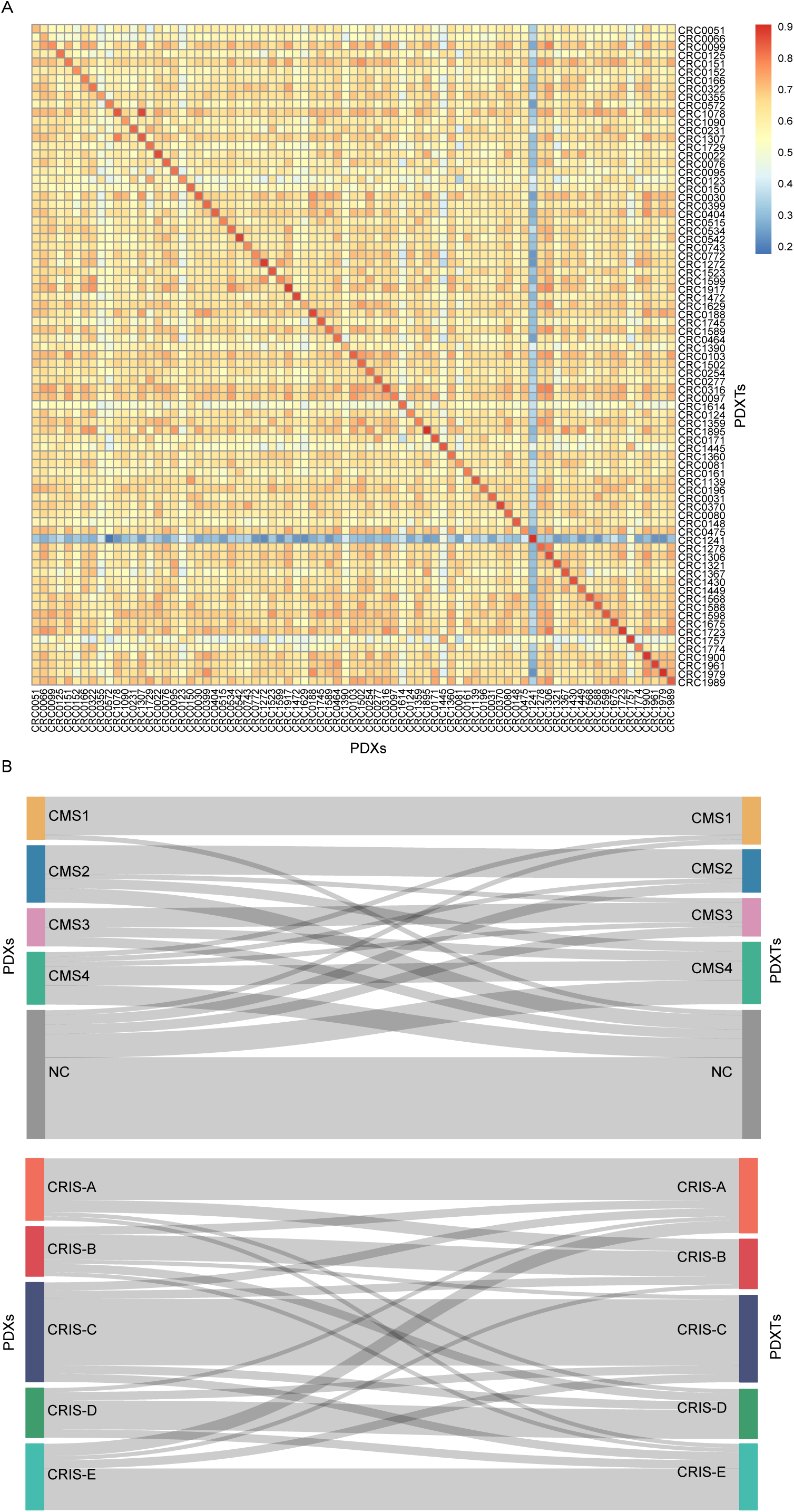
Comparative gene expression profiles and transcriptional subtype assignment in paired PDXTs and PDXs. **A,** Pearson correlations of gene expression profiles in 79 pairs of matched PDXs and PDXTs. Matched pairs, average Pearson coefficient, 0.82; unmatched pairs (*N* = 6,162), average Pearson coefficient, 0.61; matched versus unmatched pairs, *P* = 3.477e-52 by two-tailed Mann-Whitney test. **B,** CMS and CRIS subtype assignment in 79 pairs of matched PDXTs and PDXs. Consistency index between PDXs and PDXTs: CMS1, 0.69; CMS2, 0.48; CMS3, 0.40; CMS4, 0.18; non-classified, 0.45; CRIS-A, 0.43; CRIS-B, 0.33; CRIS-C, 0.55; CRIS-D, 0.41, CRIS-E, 0.45. NC, non-classified.

Gene expression profiling has been recently deployed to develop CRC classifiers with prognostic and predictive significance. The Consensus Molecular Subtypes (CMS) classifier was built on whole-tumor transcriptomes (including cancer cell and stromal/immune transcripts)^36^, whereas the CRC Intrinsic Signature (CRIS) classifier utilized PDX gene expression datasets to derive cancer cell-specific subtypes^32^. We first assigned each PDXT and PDX of the ‘super-matched’ 79 pairs to a CMS or CRIS subtype. Plausibly, many models failed CMS categorization due to lack of human stroma, which greatly contributes to CMS subtype assignment^31,32,37^; conversely, all models received a CRIS designation (Fig. 4B). We then evaluated the consistency in subtype assignment using a tailored version of the Jaccard index, whereby the number of models with the same subtype in matched PDXs and PDXTs was divided by the total number of models assigned to that subtype. This index revealed good overall correspondence, with average values across subtypes of 0.44 for both CMS and CRIS. At the level of individual subtypes, a general stability in class assignment was observed with the exception of CMS4 (Fig. 4B). This is expected, as CMS4 characteristics are dominantly driven by human stromal transcripts that are absent in XENTURION models. PDXTs therefore display representative gene expression profiles that define their identity with parental PDXs and allow their classification into RNA expression-based CRC subtypes without a substantial culture bias.

### PDXT sensitivity to cetuximab is concordant with PDX response in a large-scale population trial

The anti-EGFR antibody cetuximab is a standard-of-care treatment with demonstrated clinical benefit in patients with inoperable RAS/RAF wild-type metastatic CRC^38^. We and others have used PDX-based resources to identify determinants of responsiveness and resistance to cetuximab and to nominate novel druggable targets for cetuximab-resistant tumors^4,39–43^. As a consequence, a large part of XENTURION’s PDXTs were derived from PDX models for which annotation of sensitivity to cetuximab was available. We leveraged this information to investigate how and to what extent PDXTs may act as functional *ex vivo* surrogates of therapeutic profiles in xenografts.

Sensitivity to cetuximab was surveyed in 119 PDXTs plated at three different cell densities (1,250, 5,000 and 20,000 cells/well) in a 96-well format and cultured for one week with or without the antibody in the absence of EGF (which competes with cetuximab for receptor binding). Response was determined by measuring the ratio between treated and untreated cells and using as readouts endpoint luminescent ATP content and longitudinal cell imaging, for a total of 6,174 measurements. A coefficient of variation was calculated to evaluate consistency between biological triplicates (see Methods); by applying this filter, 116 PDXT models were selected for further analyses.

The experimental setting was overall robust and reproducible, as documented by the significant correlations between luminescence-based and imaging-based detection for all cell plating densities [Pearson coefficient, 0.52 (1,250 cells); 0.65 (5,000 cells); 0.77 (20,000 cells)], and resulted in a graded distribution of responsiveness to cetuximab treatment (Fig. 5A). We then used linear regression models to compare all these PDXT measurements with the *in vivo* tumor response (defined as the relative volume change after three weeks of treatment) in 79 matched PDXs (Supplementary Table 6). Also in this case, correlations were all positive and significant, with ATP values for the 5,000 cell-plating density showing the best performance (Pearson coefficient, 0.56) (Fig. 5B and Supplementary Fig. 9).

**Figure 5.**
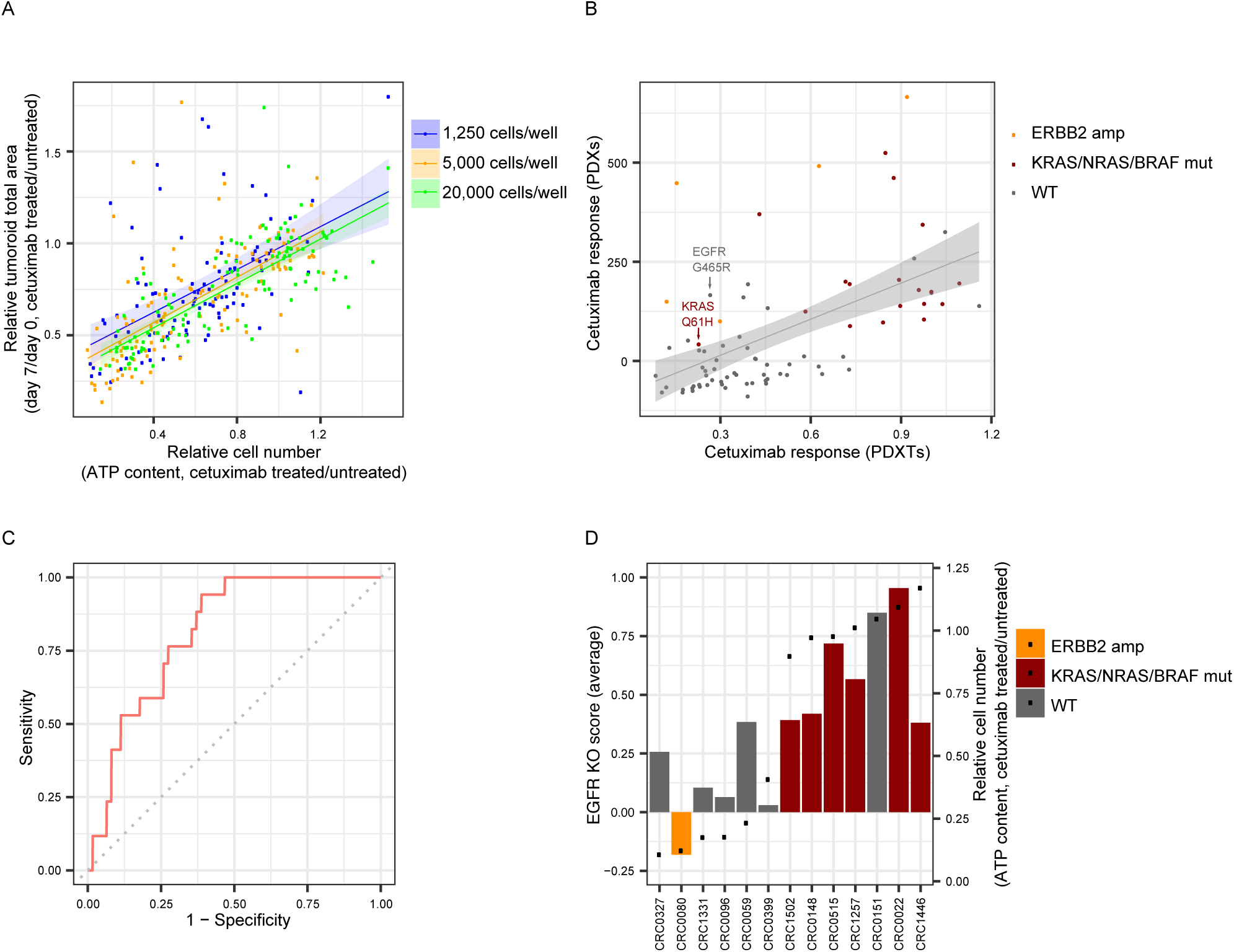
Comparative annotation of cetuximab response profiles in paired PDXTs and PDXs. **A,** Correlation of cetuximab response between values of endpoint ATP content (relative cell number) and values obtained by longitudinal cell imaging (relative tumoroid total area) in 116 PDXTs plated at different cell densities and treated with cetuximab (20 µg/mL) for one week. Each dot represents one single experiment performed in biological triplicate. Responses were assessed in 116 models for 5,000 and 20,000 cells/well, and 102 models for 1,250 cells/well. Pearson coefficient, 0.52 (*P* = 1.7e-8) for 1,250 cells; Pearson coefficient, 0.65 (*P* = 1.4e-15) for 5,000 cells; Pearson coefficient, 0.77 (*P* = 5.5e-24) for 20,000 cells. **B,** Correlation of cetuximab response in 79 pairs of matched PDXTs and PDXs. Response in PDXTs was evaluated as the ratio of viable cells after one week of treatment (20 µg/mL cetuximab, 5,000 cells/well in a 96-well format) to untreated controls; response in matched PDXs was evaluated as the percentage of tumor volume variation after three weeks of treatment (20 mg/kg, intraperitoneal injection twice a week) compared with tumor volume the day before treatment initiation. Pearson coefficient, 0.56 (*P* = 9.9e-8). **C,** ROC curve showing the performance of PDXT-based results in predicting cetuximab response *in vivo* in 79 pairs. AUC, 0.81; responders (target prediction), 17; non­responders, 62. **D,** EGFR KO scores for 13 PDXTs, distributed according to cetuximab sensitivity (black dots). Results are the average of the mean effect size of two sgRNAs against EGFR in two independent experiments, each performed in biological triplicates (with the exception of CRC0148, which was tested in three independent experiments). Pearson coefficient, 0.78, *P* = 0.0016. KO, knockout; WT, wild-type. Amp, amplification; mut, mutation; WT, wild-type.

Next, we expanded our analyses with a punctual assessment of genetic biomarkers known to confer resistance to cetuximab in patients. In line with clinical observations, tumors harboring *KRAS*, *NRAS* or *BRAF* mutations were generally refractory to EGFR blockade in both platforms (15 models out of 18 had a luminescence ratio > 0.7 when tested as PDXTs and a growth increase of more than 70% when tested as PDXs) (Fig. 5B). However, we also found some discrepant examples. First, one *KRAS* Q61H mutant model that had been categorized as a mild non-responder *in vivo* (relative tumor volume increase after three weeks of treatment, 42.35%) proved to be sensitive in the corresponding PDXTs (treated/untreated luminescence ratio, 0.23) (Fig. 5B). Interestingly, heterogeneous responses of *KRAS* Q61H mutant metastatic CRC tumors to anti-EGFR antibodies have also been observed in patients^44,45^. Second, a model with a subclonal cetuximab resistance mutation in the originating tumor (*EGFR* G465R, variant allele frequency 0.195) showed overt resistance in PDXs (relative tumor volume increase after three weeks of treatment, 166.5%) but appreciable sensitivity in the matched PDXT (treated/untreated luminescence ratio < 0.3) (Fig. 5B). In this case, the disconnect between PDX and PDXT data is due to sampling bias; the PDXs used for monitoring tumor response to cetuximab *in vivo* harbored the *EGFR* G465R mutation, whereas the PDXTs used for the *ex vivo* drug screen were derived from a sibling xenograft where the alteration was absent. The final element of divergence was found for *ERBB2* amplification, which predicts poor response to EGFR inhibition in patients with metastatic CRC^46,47^. As shown previously^39,41^, *ERBB2*-amplified PDXs failed to respond to cetuximab (Fig. 5B); however, this resistant phenotype was only partially recapitulated in PDXTs, with three models out of five displaying a certain degree of sensitivity (treated/untreated luminescence ratio ≤ 0.3) (Fig. 5B). The signaling and transformation potency of HER2 in *ERBB2*-amplified tumors is tunable by EGF stimulation^48^. On this ground, we speculate that the HER2 bypass pathway that blunts response to EGFR inhibition was below threshold in some PDXTs due to lack of EGF in the culture medium (thus, tumoroids retained sensitivity to cetuximab); conversely, the widespread availability of murine EGF in PDXs stimulated HER2 signaling to an extent sufficient to impart resistance to EGFR inhibition *in vivo*.

We reasoned that results from this population trial might prove valuable to formalize the predictive accuracy of PDXTs in modeling PDX experiments. The overall area under the curve to distinguish overtly responsive PDXs (relative tumor volume shrinkage after three weeks of treatment > 50%) from those that remained stable or progressed while on treatment was 0.81 (Fig. 5C). Based on this ROC analysis, a luminescence ratio of 0.4 in PDXTs identified 94% of matched PDXs that responded to treatment with tumor regression (Supplementary Table 6). However, the positive predictive value of pharmacologic assays in PDXTs was relatively low (FDR = 0.6), confirming the importance of model-matched *in vivo* validation during the preclinical phases of drug development. These considerations illustrate the merit of assessing the efficacy of a specific drug in a vast collection of tumoroids to advise the rational selection of models for PDX experiments.

### Gene editing recapitulates the outcome of pharmacologic target inhibition in PDXTs

Cancer dependency maps, obtained by perturbing genes with RNA interference or gene editing technologies, have provided a catalog of tumor vulnerabilities with potential clinical actionability^49,50^. These efforts have been traditionally pursued in immortalized cancer cell lines, but there is now increasing recognition that functional genomics screens in tumoroids would be better representative of cancer biology and diversity^51^. With this in mind, we sought to explore whether genetic versus pharmacologic inhibition of an index cancer dependency gene results in similar or different effects on PDXT viability. Given the large number of models with known response to cetuximab, EGFR was selected as a target, and CRISPR-Cas9 technology was applied to systematically disrupt the *EGFR* gene in 13 representative PDXTs with variable sensitivity to cetuximab (Supplementary Fig. 10A and B).

Two different sgRNAs targeting EGFR in exon three were independently transduced into Cas9-expressing PDXTs. Seven days after infection, tumoroids were processed for luminescence-based detection of ATP content. A knockout (KO) score was calculated by intra-model normalization of the viability outputs of EGFR-edited PDXTs to conditions of negligible influence on cell fitness (deletion of a neutral/non-essential gene) or strong influence (deletion of a lethal/essential gene) (see Methods). The consequences of EGFR deletion on PDXT viability were similar for the two sgRNAs (Pearson coefficient, 0.84) (Supplementary Fig. 10C), supporting robustness and reproducibility of the dataset. Remarkably, the overall outcome of EGFR genetic ablation was significantly correlated to that of cetuximab treatment (Pearson coefficient, 0.78), and in some cases a direct quantitative correspondence between the extent of pharmacologic sensitivity and the impact of gene deletion could be observed (Fig. 5D); for example, a RAS wild-type model that proved to be highly refractory to cetuximab treatment was also poorly impacted by EGFR disruption (CRC0151); in a complementary fashion, EGFR KO was severely detrimental in an *ERBB2*-amplified PDXT that was also particularly sensitive to cetuximab (CRC0080) (Fig. 5D). Collectively, these findings underscore the power and reliability of using genetic approaches in tumoroids for preclinical characterization of drug targets and to interrogate the effects of loss-of-function alterations in cancer-relevant genes.

### An in silico, *ex vivo* and *in vivo* funneling approach identifies actionable co-dependencies that attenuate response to cetuximab

The concordance of molecular profiles and therapeutic annotation in matched PDXTs and PDXs prompted us to embark on a discovery effort aimed to identify adaptive dependencies in models that were sensitive to, but not eradicated by, EGFR inhibition. To do so, we followed a principled approach meant to triage candidate vulnerabilities using sequential selection bottlenecks, with the final aim to nominate only those targets that passed strict validation criteria.

We started by analyzing transcriptional responses to drug pressure in cetuximab-sensitive models (33 PDXs and 12 PDXTs) treated with the antibody, with the assumption that some upregulated gene products may adaptively convey compensatory signals to contrast EGFR inhibition. Similar to basal (pre-treatment) profiles, also on-treatment gene expression changes were coherent in PDXTs and PDXs (Pearson coefficient, 0.8) (Fig. 6A). The list of genes that were upregulated by cetuximab in all PDXs and PDXTs examined was refined by removing low-expressed ones and those that were also modulated by treatment in an independent set of 21 cetuximab-resistant PDXs. The resulting compendium of 1,916 genes was restricted to a subset of 119 genes encoding druggable targets, based on a two-tiered selection strategy: i) a target tractability assessment using published criteria^50^, such as the availability of compounds in clinical or preclinical development and/or the documentation of an associated response biomarker; ii) data output from the Drug-Gene Interaction Database (www.dgidb.org), a web resource that provides information on drug-gene interactions and druggable genes. The 119 gene subset was further narrowed down to 13 candidates through a shortlisting process that considered target conceptual novelty and translational potential as well as known potency and *in vivo* bioavailability of the corresponding specific drugs (Supplementary Fig. 11 and Supplementary Table 7).

**Figure 6.**
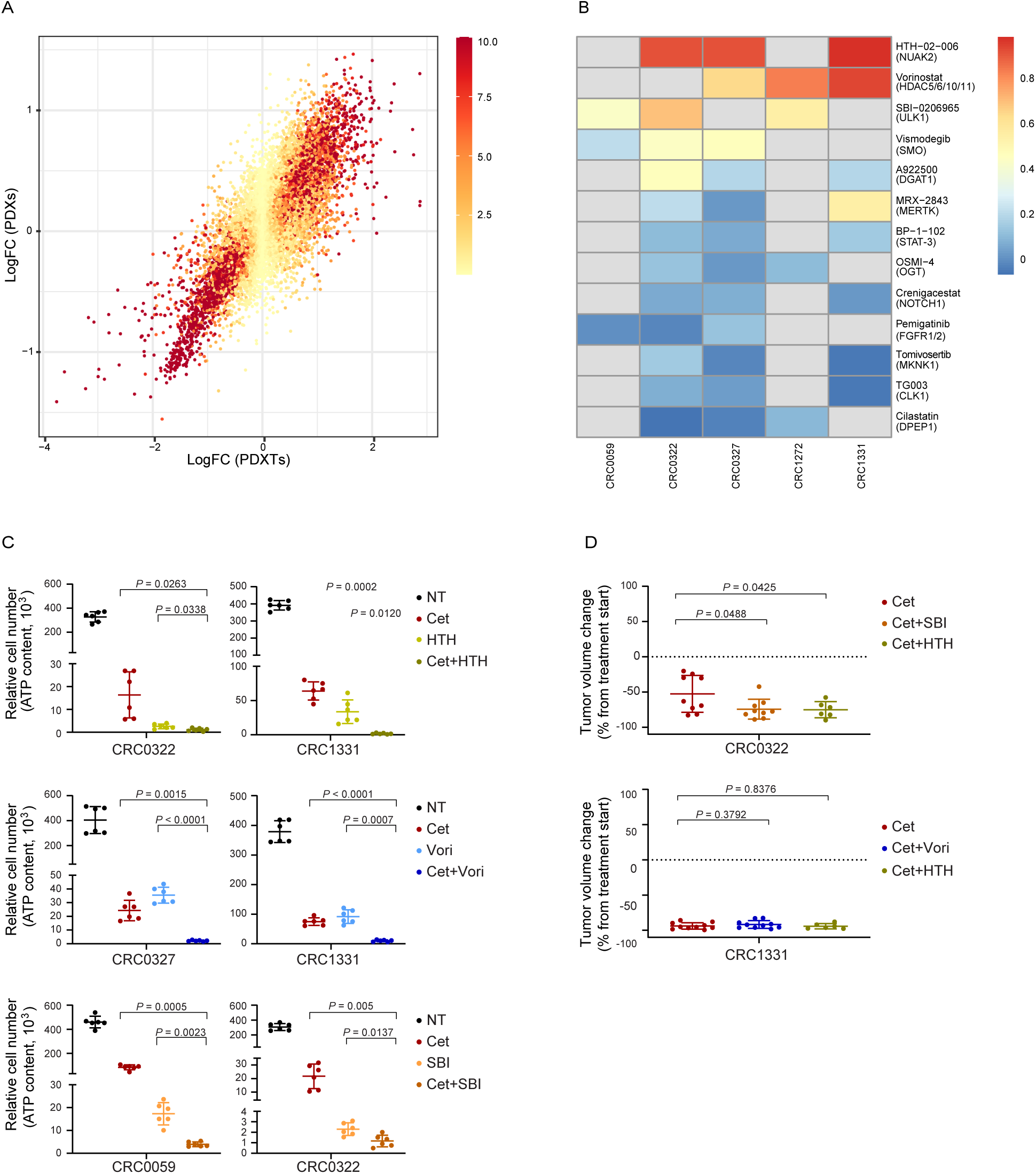
Drug screen in PDXTs and therapeutic validation trial in PDXs. **A,** Correlation of gene expression changes between 33 PDXs exposed to cetuximab for three days (*N* = 28 models) or six weeks (*N* = 6 models) and 12 PDXTs exposed to cetuximab for three days. The scatterplot shows log-fold changes between treated and untreated samples for 19,716 genes. Color shading reflects differential expression *P* values, as assessed by DESeq2. Pearson coefficient, 0.8 (*P* < 2.2e-308). **B,** Maximum inhibition scores for 13 candidate drugs, each tested in three different PDXT models. PDXTs were treated for 48 hours (5,000 cells/well in a 96-well format), and viability was assessed by endpoint ATP content. For each drug, three independent experiments, each in biological triplicates, were performed. Maximum inhibition score was calculated as the difference between the viability of untreated cells and that recorded at maximum drug dosage, normalized against the viability of untreated cells and averaged for results of the three independent experiments. Maximum drug dosage: HTH-02-006, 16 µM; vorinostat, 10 µM; SBI-0206965, 20 µM; vismodegib, 50 µM; A922500, 100 µM; MRX-2483, 1 µM; BP-1-102, 1 µM; OSMI-4, 20 µM; crenigacestat, 5 µM; pemigatinib, 1 µM; tomivosertib, 5 µM; TG003, 50 µM; cilastatin, 0.5 µM. Drug targets are specified. **C,** Relative cell viability (ATP content) in PDXT models treated for one week (CRC0327 e CRC1331, 5,000 cells/well in a 96-well format) or three weeks (CRC0059 and CRC0322, 1,000 cells/well in a 96-well format) with SBI-0206965 (10 µM), HTH-02-006 (4 µM) and vorinostat (1.25 µM), either as monotherapies or in combination with cetuximab (20 µg/ml for one-week treatments and 5 µg/ml for three-week treatments). Results represent the means ± SD of two independent experiments performed in biological triplicates (*N* = 6). Statistical analysis by Brown-Forsythe and Welch ANOVA followed by Dunnett’s T3 multiple comparison test. **D,** Tumor volume changes in PDXs from mice treated with the indicated modalities for 4 weeks. Cetuximab, 20 mg/kg (intraperitoneal injection twice a week); HTH-02-006, 10 mg/kg (intraperitoneal injection twice a day); SBI-0206965, 20 mg/kg (intraperitoneal injection three times a week); vorinostat, 50 mg/kg (intraperitoneal injection three times a week). Dots represent volume changes of PDXs from individual mice, and plots show the means ± SD for each treatment arm. *N* = 6 to 10 animals per each treatment arm. Tumor volume changes of the placebo arm are shown in Supplementary Fig.13. Statistical analysis by two-tailed unpaired t test with Welch’s correction. Cet, cetuximab; HTH, HTH-02-006; SBI, SBI-0206965; Vori, vorinostat.

To preliminarily assess whether the prioritized hits were valuable therapeutic targets, we performed short-term (48h) viability assays using well-characterized chemical inhibitors of the 13 candidates. Each inhibitor was tested in three PDXTs in the absence or presence of cetuximab, with model selection guided by robust cetuximab-induced overexpression of the targeted gene (in most cases, one or more models were used to test more than one drug) (Supplementary Table 8). The compounds were evaluated in a four-point dose-response assay with biological triplicates and in three independent experiments, using the conditions that proved to be the most accurate in recapitulating response *in vivo* (5,000 cells/well in a 96-well format and luminescence-based detection of ATP content as viability readout), for a total of 3,510 measurements (Supplementary Fig. 12 and Supplementary Table 9). Drug doses were calibrated using literature data and publicly available pharmacologic profiles^52^, with a maximum dose for each inhibitor equal to twice the standard IC_50_ value. From this first set of assays, three compounds stood out for having the highest maximum inhibition score as single agents (*i.e.*, their maximum dose had the strongest impact on cell viability) (Fig. 6B and Supplementary Table 9): HTH-02-006, targeting NUAK, a member of the AMPK subfamily of serine/threonine protein kinases; vorinostat, targeting histone deacetylases (HDACs); and SBI-0206965, targeting the serine/threonine kinase ULK1. Interestingly, these targets are variably involved in promoting autophagy, a homeostatic mechanism triggered by cellular stress^53^.

Blockade of the 13 candidates together with cetuximab incrementally reduced cell viability, again with HTH-02-006, vorinostat and SBI-0206965 showing the highest maximum inhibition scores (Supplementary Fig. 12 and Supplementary Table 9). We then explored the interaction between cetuximab and the three top inhibitors under conditions of longer drug exposure to favor the implementation of adaptive reactions over time. In this setting, combination therapy with cetuximab and HTH-02-006, vorinostat or SBI-0206965 was significantly more effective than single-agent treatments and outperformed cetuximab alone (Fig. 6C).

Finally, pharmacologic experiments were translated into the *in vivo* setting. PDX model CRC0322 was chosen for administration of HTH-02-006 and SBI-0206965 because the corresponding PDXT was particularly sensitive to both drugs; following a similar reasoning, PDX model CRC1331 was chosen because the matched PDXT displayed the highest maximum inhibition score for both vorinostat and HTH-02-006 (Fig. 6B). As expected, both PDX models were strong responders to cetuximab monotherapy (Fig. 6D). While single agent-treatment with HTH-02-006, vorinostat or SBI-0206965 had negligible or null effect on tumor growth (Supplementary Fig. 13), the combination of HTH-02-006 or SBI-0206965 with cetuximab proved to be more effective than cetuximab alone in reducing tumor size in CRC0322 PDXs (Fig. 6D). In CRC1331, combination therapy of cetuximab with either vorinostat or HTH-02-006 did not significantly outperform cetuximab monotherapy (Fig 6D); however, it is worth noting that response to cetuximab was exceptionally profound in this PDX (mean tumor volume decrease, 93.77%), which may have masked the contribution of the other drug to tumor regression. In essence, these results emphasize the value of integrative molecular and biological data, coupled with vast availability of experimental models, for ‘aggressive’ prioritization of targets with meaningful translational potential.

## DISCUSSION

XENTURION is a living biobank of matched xenografts and tumoroids from patients with metastatic CRC, with associated multi-dimensional molecular data and biological annotation. A similar resource has been recently developed for treatment-refractory and metastatic breast cancers^54^. Models in XENTURION recapitulated the genetic heterogeneity of metastatic CRC – as evidenced by a representation of somatic variants and copy number alterations analogous to that found in clinical datasets of CRC metastases – and showed high intra-pair genetic and transcriptional similarity. We have not thoroughly analyzed whether this concordance also extends to parental tumors in donor patients, and we do not exclude that some patient-derived models might be divergent from originating samples due to selection bottlenecks and/or evolutionary drift. However, we believe that the molecular makeups displayed by individual models well reflect those of spontaneous tumors in the general population, and we regard our resource as an illustrative census of metastatic CRC as a whole. This is even more so when considering that our study was not designed to prospectively guide human care in donor patients, an approach that would require tumors in patients to have perfect equivalents in culture and mice for dependable returning of results to the clinic^55^.

It is commonly assumed that tumoroids may be used to shortlist drug candidates before *in vivo* validation, but their reliability in anticipating xenograft results remains to be established^19^. The vast number of matched PDXT-PDX pairs developed in XENTURION enabled performing correlations with an adequate statistical power, which facilitated the formal assessment of the accuracy of PDXTs in predicting cetuximab response in the corresponding PDXs. To our knowledge, this is the first large-scale drug efficacy study in which a systematic comparison of therapeutic response between PDXT-PDX pairs was attempted, and the first report gauging the diagnostic ability of PDXTs in informing subsequent steps of *in vivo* experimentation. We also found that the effect of pharmacologic versus genetic inactivation of EGFR on tumoroid viability was similar. Therefore, mapping cancer dependencies by methodical gene knockout in tumoroids is expected to deliver results comparable to those achieved by drug-mediated perturbation; this should help prioritize pharmaceutical development pipelines for those drug-orphan targets that, when deleted, drastically impair tumoroid growth^51^.

Our PDXT pharmacologic characterization uncovered experimental drugs with therapeutic potential. We reasoned that some cetuximab-induced genes might contribute to adaptive tolerance to EGFR inhibition, and blockade of their protein products should enhance cetuximab sensitivity. With this premise, genes concordantly and specifically upregulated by cetuximab in PDXs and PDXTs were prioritized based on tractability. Potential targets were further selected according to the extent to which their inhibition impacted tumoroid viability, and based on this ranking three candidates (NUAK2, ULK1, and HDACs) were shortlisted. Accordingly, their neutralization deepened response to cetuximab. Notably, all three targets are involved in autophagy, a stress-induced survival pathway during which vesicles called autophagosomes engulf damaged organelles and unfolded proteins and deliver them to the lysosome for degradation^53^. In particular, NUAK2 and ULK1 participate in the formation of autophagic vesicles^53,56^, and HDACs control the fusion of autophagosomes to lysosomes^57^. Although we have not formally documented induction of autophagy after cetuximab treatment, it is tempting to speculate that autophagy may be an alternate survival strategy deployed by CRC tumors to counteract the growth-inhibitory effect of EGFR inactivation, and autophagy inhibitors may cooperate with cetuximab to achieve deeper tumor regressions.

Results gathered from our experiments also highlight the limitations of XENTURION. The minimal composition of the growth medium (which contained EGF as the sole growth factor) was instrumental to standardizing culture conditions and sufficient to ensure the growth of most tumoroids, but favored the establishment of EGF-dependent models – such as those derived from metastases of left-colon primary tumors - and disfavored that of models with a higher reliance on WNT signals - namely, those with β-catenin rather than APC mutations. In the same vein, viability assays with cetuximab were biased by the need to remove EGF from the medium to avoid antibody competition for the same receptor binding sites. This procedure was conducive to some ‘false positives’ in PDXTs, as exemplified by the observed sensitivity of *ERBB2*-amplified tumoroids to cetuximab, which was not confirmed in matched PDXs. In this case, it is arguable that the poor culture milieu experienced by PDXTs made them more susceptible to EGFR inhibition than corresponding xenografts in mice, which likely were exposed to survival signals conveyed by the host. More in general, the influence of culture media composition on tumoroid ability to predict therapeutic responses is substantial^58^, and additional work is needed to define cocktails of growth factors with a proper balance between the necessity of methodological synthesis and that of coping with idiosyncratic model characteristics.

We have implemented protocols for sharing XENTURION models and associated data responsibly with the scientific community, while protecting patient confidentiality and the right to withdraw informed consent. We have also planned the development of a user-friendly web portal that will provide open access to the molecular and biological characteristics of the models and visualization tools. In conclusion, XENTURION is a rich resource that augments and integrates existing genomic, transcriptomic and functional datasets of CRC tumoroid collections. Being open to external parties, this resource is expected to yield fertile ground for catalyzing independent translational exploration and fostering new discoveries.

## METHODS

### Specimen Collection and Annotation

Tumor samples were obtained from donor patients treated by liver metastasectomy at the Candiolo Cancer Institute (Candiolo, Torino, Italy), Ospedale Mauriziano Umberto I (Torino), Città della Salute e della Scienza di Torino – Presidio Molinette (Torino), and Grande Ospedale Metropolitano Niguarda (Milano, Italy). All patients provided informed consent. Samples were procured and the study was conducted under the approval of the Review Boards of the Institutions (PROFILING protocol No. 001-IRCC-00IIS-10). Clinical and pathological data were entered and maintained in our prospective database. The deidentified clinical information in this study is published in accordance to the ethics guidelines of the PROFILING protocol.

### Overview of the sequenced samples

Targeted DNA sequencing was performed in 147 tumoroids; 149 PDXs; 4 human samples; and 1 control sample with known mutational status of the covered genes (Supplementary Table 10); 129 of the sequenced tumoroids ended the validation process successfully. Shallow sequencing was performed in 146 tumoroids; 149 PDXs; 4 human samples; and 3 control samples with known copy number profiles (Supplementary Table 10); 129 of the sequenced tumoroids ended the validation process successfully. RNA sequencing was performed in 220 tumoroids (all passed quality checks); 480 PDXs (470 passed quality checks); 68 human samples (65 passed quality checks); 31 pairs of cetuximab-treated and untreated PDXTs (all passed quality checks); 43 PDXs exposed to cetuximab for three days (all passed quality checks) and 42 untreated PDX controls (all but one passed quality checks); 52 PDXs exposed to cetuximab for six weeks (all but one passed quality checks) and 48 untreated PDX controls (all but one passed quality checks) (Supplementary Table 11).

### DNA and RNA extraction, sgRNA PCR amplification and Sanger sequencing

Total DNA and RNA were extracted using the Maxwell® Instrument (Promega) following manufacturer’s instructions. EGFR-amplified products were obtained by PCR using the following primers: 5’-CAGGAGGTGGCTGGTTATGT-3’ and 5’-TTCTCCGAGGTGGAATTGAG-3’. Sanger sequencing was performed by using the following primer: 5’-TTCTCCGAGGTGGAATTGAG-3’. To obtain bulk RNA-seq data, RNA was extracted using miRNeasy Mini Kit (Qiagen), according to the manufacturer’s protocol. The quantification and quality analysis of RNA was performed on a Bioanalyzer 2100 (Agilent), using RNA 6000 Nano Kit (Agilent).

### Analysis of mutant variants

Targeted sequencing was performed on an Illumina NovaSeq 6000 instrument by IntegraGen SA. Paired-end 2×100bp reads were obtained, aiming at ∼500x depth for each sample on the targeted regions (402.4 kbp). The panel (Twist Bioscience) covered exons of 116 genes (Supplementary Table 12) from a manually curated list of recurrently mutated drivers in CRC. Initial quality check (QC) was performed with FastQC (v0.11.9) by IntegraGen; for PDX samples, murine reads were filtered using Xenome (v1.0.0) with default parameters, after building k-mer indexes for the human genome (GRCh38, downloaded from https://gdc.cancer.gov/about-data/gdc-data-processing/gdc-reference-files) and murine genome (mm10, obtained from iGenomes). The GATK Best Practices Workflow^59^ for somatic mutation calling was followed to perform mutation calling with Mutect2 (bwa v0.7.17-r1188 - parameters -K 100000000 -Y, GATK 4.1.4.0); alignment was performed versus GRCh38; alignment metrics were gathered with picard CollectHsMetrics (Supplementary Table 13). dbSNP (for quality recalibration, downloaded from NCBI, ftp://ftp.ncbi.nih.gov/snp/organisms/human_9606/VCF/All_20180418.vcf.gz) and Gnomad (release 2.1.1) were used as external references for common human polymorphisms. The resulting list of filtered mono-allelic mutations was annotated using PCGR (version 0.9.1)^60^; to avoid germline contamination, only coding mutations found in tiers ≤ 3 (https://pcgr.readthedocs.io/en/latest/tier_systems.html) were kept for further analyses. TCGA and MSK-IMPACT mutational data was downloaded from (https://www.cbioportal.org/study/clinicalData?id=crc_msk_2017 and https://www.cbioportal.org/study/summary?id=coadread_tcga_pan_can_atlas_2018-Mutated_Genes.txt). To compute frequencies of clonal events only mutations with AF > 0.2 were kept. To compare mutational landscapes between PDXs and PDXTs a more lenient filter (AF > 0.05) was applied.

### Copy number profiles

Shallow-sequencing was performed on Illumina NovaSeq S4 instrument by Biodiversa SRL (Supplementary Table 14). Paired-end 2×150bp reads were obtained, aiming at 0.65x depth for each sample. Initial QC was performed with FastQC (v0.11.8); for PDX samples, murine reads were filtered using Xenome (v1.0.0) with the same procedure described for targeted sequencing. Total or human classified reads were aligned versus GRCh38 with bwa (v0.7.17-r1188 - parameters -K 100000000 -Y) following GATK best practices; then, duplicates were marked using picard (2.18.25). Alignment metrics were gathered with samtools (v1.9) flagstat and picard CollectWgsMetrics (Supplementary Table 15). Segmented fold changes were obtained with QDNAseq (v1.22.0)^61^ with a bin size of 15kb (annotations obtained from QDNAseq.hg38), using default parameters and pairedEnds=TRUE. Before computing correlations, a pseudocount of 1 was added to all fold changes to compute log2; a pseudocount of 0.01 was used for visualization. Log2 values with a pseudocount of 1 in .seg format were used as input for the GISTIC2.0 module in GenePattern (https://cloud.genepattern.org, default parameters and TRHuman_Hg38.UCSC.add_miR.160920.refgene.mat, Gistic v2.0.23). TCGA Gistic output (specifically all_thresholded.by_genes.txt) was downloaded from the GDCl (https://gdc.cancer.gov/about-data/publications/coadread_2012); for MSK-IMPACT, the file with segmented signals was downloaded from cBioPortal (https://www.cbioportal.org/study/clinicalData?id=crc_msk_2017) and GISTIC was run as previously described.

### Gene expression analyses

#### Quality control checkpoints

Total RNA was processed for RNA-seq analysis with the TruSeq RNA Library Prep Kit v2 (Illumina) following manufacturer’s instructions, and sequenced on a NextSeq 500 system (Illumina) by Biodiversa SRL. Single-end 151bp reads were obtained, aiming at 20M reads for each sample. Read counts were obtained using an automated pipeline (https://github.com/molinerisLab/StromaDistiller), that uses a hybrid genome composed of both human and mouse sequences to exploit the aligner ability to distinguish between human-derived reads, representing the tumor component, and mouse-derived reads, representing the murine host contaminating RNA material. Reads were aligned using STAR^62^ (version 2.7.1a, parameters --m outSAMunmapped Within --outFilterMultimapNmax 10 -- outFilterMultimapScoreRange 3 --outFilterMismatchNmax 999 --outFilterMismatchNoverLmax 0.04) versus this hybrid genome (GRCh38.p10 plus GRCm38.p5hg38 with Gencode version 27 and mouse GRCm38 with Gencode version 16, indexed with standard parameters and including annotation information from the GENCODE 27 plus m16 comprehensive annotation). Aligned reads were sorted using sambamba^63^ (version 0.6.6) and only non-ribosomal reads were retained using split_bam.py^64^ (version 2.6.4) and rRNA coordinates obtained from the Gencode annotation and RepeatMasker track downloaded from UCSC genome browser hg38 and mm10. featureCounts^65^ (version 1.6.3) was run with the appropriate strandness parameter (-s 2) to count the non multi-mapping reads falling on exons and reporting gene-level information (-t exon -g gene_name) using combined Gencode basic gene annotation (27 plus m16).

Sequencing data was available for 1015 samples, but different filtering criteria resulted in 999 samples surviving quality checks. These criteria included: i) number of total reads ≥ 15M; ii) reads assigned to genes by feature counts ≥ 60%; iii) reads assigned to human genes over the total of assigned reads ≥ 30%. By applying these filters, only samples with at least 5M human reads were considered for analysis. To remove samples with lymphomatous characteristics^32^, 2 criteria were applied: i) Principal Component analysis of expression data (samples with PC2 ≥ 30 were discarded): ii) computation of a sample-level score for a leukocyte expression signature^31^, averaging FPKM values for all the signature genes (samples with an average leukocyte signature ≥ 48 were discarded). Positivity for either criterion flagged samples as lymphomatous and excluded them from analysis. The overall concordance of the PC score with the leukocyte expression signature is shown in Supplementary Fig. 14.

Samples were sequenced in five different batches. This did not largely affect sample separation based on Principal Component analysis (Supplementary Figure 14). However, when possible, batch correction was applied in downstream differential analyses. Variance stabilized expression and robust FPKM values were obtained using DESeq2^66^ (version 1.26.0), tmm using edgeR (version 3.28.1) starting from the read counts assigned to human genes only.

#### Differential expression analysis

Differentially expressed genes (DEGs) between original tumors, PDXs and PDXTs were obtained using R package DESeq2 (v1.26.0) with the formula “∼batch + type + sample”. By applying this formula, ‘batch’ was used to correct for sequencing batches; ‘type’ specified sample origin (surgical specimen, PDX, or tumoroid); and ‘sample’ was an identification tag assigned to each original tumor (sample was added in the formula to obtain DEGs between sample types, taking into account tumor identity). Genes with more than 5 reads in only 1 sample were removed before testing for differential expression. DEGs were identified using |LFC| ≥ 0.5849625 and adjusted *P* values < 0.05. Samples were selected starting from PDXTs that had passed the validation process and their paired PDXs and HLMs. The obtained DEGs were used to perform GO enrichment analysis with R libraries ClusterProfiler^67,68^ (v3.14.3), DOSE^69^ (v3.12.0), msigdbr^70^ (v7.4.1) and enrichplot (v1.6.1). Graphical representations of GO Biological Process enrichments were obtained with REVIGO (http://revigo.irb.hr/)^71^, which was manually run online on 05/12/22 with the list of GO terms with adjusted *P* values < 0.05 as input and selecting ‘small’ as output size. The final plots were obtained by manual highlighting of a subset of relevant terms with ggplot.

DEGs between untreated and cetuximab-treated samples were obtained using the formula “∼sample + treat” for PDXTs and “∼sample + time + treat” for PDXs; ‘treat’ indicated if sample was treated or untreated; ‘time’ indicated time of PDX exposure to treatment (three days or six weeks). The list of significant DEGs in PDXTs was refined by successive filters (Supplementary Figure 11 and Supplementary Table 7), considering only: i) genes upregulated by cetuximab in PDXTs (log2fold change > 0); ii) genes also upregulated by cetuximab in responsive PDXs (log2fold change > 0 and adjusted *P* value < 0.05); iii) genes with no significant expression changes (adjusted *P* value ≥ 0.05) after exposure of cetuximab-resistant PDXs to antibody treatment; iv) genes with a baseMean, reported by DESeq2, > than their overall median. The list of resulting gene symbols was annotated for tractability considering buckets 1-7 in Behan et al.^50^ and used as input in the Search Interaction form of the Drug-Gene Interaction Database (www.dgidb.org) on 26/03/2021.

#### Transcriptional subtype assignment

CMS subtypes were called on the vsd matrix for PDXTs, PDXs and HLMs separately, using R package CMScaller^72^ (v2.0.1) with parameters FDR = 0.05 and RNAseq = TRUE. CRIS subtypes were also called on the vsd matrix with the R package CRISclassifier^32^ (v1.0.0).

#### Hierarchical clustering

Hierarchical clustering (method = ‘complete’, distance metric 1 - Pearson correlation) was performed on vsd expression data, keeping replicate samples as separate entities. Filters were applied to eliminate low expressed genes and to maintain those with highly variable expression; specifically, genes with average expression across samples larger than the overall median of averages and the top 10% genes with higher standard deviation were analyzed.

#### Inference of tumor’s primary tissue of origin

Inference of tumor’s primary tissue of origin was obtained with CUP-AI-Dx (https://github.com/TheJacksonLaboratory/CUP-AI-Dx, docker yuz12012/ai4cancer:product)^35^ with log2 tmm with a pseudocount of 1 as expression values.

### Patient-derived tumoroid cultures

Tumoroids of CRC liver metastases were established from PDX explants or directly from surgical resections of donor patients. Tumor specimens (0,5cm x 0,5cm) were chopped with a scalpel and washed with PBS. After centrifugation, the final cell preparation was embedded in Matrigel® (Corning) or Cultrex Basement Membrane Extract (BME Type II or Ultimatrix RGF BME, R&D Systems) and dispensed onto 24-well plates (Corning). After 10-20 minutes at 37°C, culture medium was added. Complete tumoroid medium composition was the following: Dulbecco’s modified Eagle medium/F12 supplemented with penicillin-streptomycin, 2 mM L-glutamine, 1mM n-Acetyl Cysteine, B27 (Thermo Fisher Scientific), N2 (Thermo Fisher Scientific) and 20 ng/ml EGF (Sigma-Aldrich). Tumoroids were tested for Mycoplasma and maintained at 37°C in a humidified atmosphere of 5% CO_2_. Periodic checks of sample identity with the original human specimen (liver metastasis and, when available, normal liver) were performed using a 24 SNP custom genotyping panel (Diatech Pharmacogenetics), and results were analyzed using the MassARRAY Analyzer 4 (SEQUENOM® Inc, California). Culture expansion and biobanking were managed using the Laboratory Assistant Suite^73^.

### Viability assays

Pharmacologic experiments were performed in 96-well plates with a thin layer of BME in each well. PDXTs were washed with PBS, incubated with trypsin-EDTA solution for 5 minutes at 37°C and vigorously pipetted to obtain a single cell suspension. Cells were seeded in 2% BME culture medium in the absence of EGF. After 1-2 days from seeding, PDXTs were treated with the modalities indicated in the figure legends. Cetuximab was provided by Merck; HTH-02-006 was purchased from Aobious; all other small-molecule inhibitors were purchased from Selleckchem. Cell viability was measured by ATP content using the Cell Titer-Glo luminescent assay kit (Promega). For cell imaging, bright-field images were acquired on a Cytation 3 cell imaging multi-mode reader (BioTek Instruments) with a 4x Olympus objective. Images were mounted by stitching together 12 different fields of view in order to completely cover each well of the plate. Images were then processed according to a custom-made Fiji script^74^ to segment tumoroid areas. Edge detection algorithms were used to get rid of the non-uniform brightness resulting from illuminating a multi-well plate, followed by a thresholding step and a morphological filling. Segmented objects were then filtered according to their size and aspect ratio and total tumoroid area was used as a proxy for growth. Data dispersion was assessed by applying a coefficient of variation (CV), calculated as the ratio of the standard deviation of untreated or treated cells to the average of untreated cells. Experiments were excluded when none of the three cell densities tested (1,250, 5,000 or 20,000 cells/well in a 96-well format) for at least one of the two readouts (ATP content and longitudinal cell imaging) displayed CV values lower than 0.25.

### CRISPR-Cas9 gene editing and dependency assays

PDXTs were transduced with a Cas9-expressing pKLV2-EF1a-BsdCas9-W lentiviral vector (Addgene) and selected with 10 µg/mL blasticidin (Thermo Fisher Scientific). sgRNAs were cloned into the pKLV2-U6gRNA5(BbsI)-PGKpuro2AZsG-W lentiviral vector (Addgene). sgRNA sequences were designed to target the following genes: i) *EGFR* (sgEGFR#1, GATAAGACTGCTAAGGCAT; sgEGFR#2, GCAAATAAAACCGGACTGA); ii) the essential gene *PLK1* (GCGGACGCGGACACCAAGG); the non­essential gene (CYP2A13, GTCACCGTGCGTGCCCCGG). After transduction with the sgRNA vectors, Cas9-expressing PDXTs were seeded at 5,000 cells/well in 96-well plates in the presence of blasticidin and 2 µg/ml puromycin (Sigma Aldrich) and cultured for seven days. Endpoint cell viability was measured using the Cell Titer-Glo luminescent assay kit (Promega) and normalized to day one. The ATP content values of PLK1-KO cells were subtracted to the ATP content values of EGFR-KO cells (difference 1) and the ATP content values of CYP2A13-KO cells (difference 2). Then, a KO score was calculated as the ratio of difference 1 to difference 2. Only PDXTs in which endpoint viability values after *PLK1* KO were at least 30% less than those of *CYP2A13* KO cells were included in the analyses.

### Western blot analysis

Four days after sgRNA transduction, PDXTs were harvested and total cellular proteins were extracted with boiling Laemmli buffer (1% SDS, 50 mM Tris-HCl pH 7.5, 150 mM NaCl). After sonication, total proteins were quantified using the BCA Protein Assay Reagent Kit (Thermo Fisher Scientific), electrophoresed on precast polyacrylamide gels (Invitrogen), and transferred onto nitrocellulose membranes using a Trans-Blot Turbo Blotting System (Biorad). Membrane-bound antibodies were detected by the enhanced chemiluminescence system (Promega). Primary antibodies were rabbit anti-EGFR (Cell Signaling Technology #4267, 1:1,000 dilution) and mouse anti-vinculin (Sigma-Aldrich, V9131, 1:2,500 dilution).

### PDX studies

Tumor implantation and expansion were performed as previously described in 6-week-old male and female NOD/SCID mice^39^. All procedures using live animals were reviewed and approved by the Candiolo Cancer Institute Institutional Animal Care and Use Committee (IACUC) and by the Italian Ministry of Health (authorization 37/2022-PR) and complied with all relevant ethical regulations. Mice with established tumors (average volume ∼300 mm^3^) were randomized and treated with the modalities indicated in the figures. Vehicle compositions were the following: 10% DMSO and 5% dextrose for HTH-02-006; 5% Tween-80 and 5% polyethylene glycol (PEG) 400 in PBS for SBI-0206965; and 25% hydroxypropyl-β-cyclodextrin, 5% DMSO, and 45% PEG 400 for vorinostat. Tumor size was evaluated once weekly by caliper measurements, and the approximate volume of the mass was calculated using the formula 4/3π ▪ (d/2)^2^ _·_ D/2, where d is the minor tumor axis and D is the major tumor axis. The maximum tumor diameter approved by the IACUC and the Italian Ministry of Health was not exceeded. Results were considered interpretable when a minimum of four mice per treatment group reached the prespecified endpoints (at least 3 weeks on therapy or development of tumors with the largest diameter ≥ 10 mm). No statistical methods were used to predetermine sample sizes; sample sizes were similar to those reported in previous publications^41,43^ and conformed to PDX minimal information standards^75^. Operators were not blinded during measurements. *In vivo* procedures, including animal randomization, and related biobanking data were managed using the Laboratory Assistant Suite^73^.

### Statistical and bioinformatics analyses

The number of biological (nontechnical) replicates for each experiment is reported in the figure legends, alongside the adopted statistical tests and metrics. For comparisons between two groups, statistical analyses were performed using two tailed *t* tests, applying Welch’s correction for unpaired tests. Similarity estimates between tumoroids and xenografts to compare matched and unmatched models were performed using two-tailed Mann-Whitney tests. For experiments with more than two groups or when comparing dose-response curves in tumoroids, one-way ANOVA was used. When not equal SDs were assumed, Brown-Forsythe and Welch ANOVA was applied. In case of multiple testing, the Dunnett’s T3 multiple comparisons test was adopted. All graphs and statistical analyses were performed usingGraphPad Prism (v9.0) and R (v3.6.3), its base packages and the following libraries: precrec (0.12.9), ggplot2 (v3.3.0), pheatmap (v1.0.12), ComplexHeatmap (v2.2.0), sjPlot (2.8.10) and circlize (v0.4.15); with the exception of the CN heatmap (Fig 3B) that was made with python3/matplotlib (v3.7.3/3.4.3) and the sankeys plot (Fig 4B), made with the package networkD3 (v0.4) and its function sankeyNetwork.

### Materials availability

PDX models and tumoroids will be made available to the community (with the exception of for-profit entities) under material transfer agreement; requests will be granted by contacting the corresponding authors.

### Data availability

All sequencing data reported here have been deposited in EGA in the following studies: EGAS00001006697 (targeted DNA sequencing); EGAS00001006696 (shallow sequencing); EGAS00001007024 (RNA sequencing of tumoroids); EGAS00001006492 (RNA sequencing of PDXs); EGAS00001006746 (RNA sequencing of original human samples); EGAS00001006601 (cetuximab-treated and untreated tumoroids); EGAS00001006973 (cetuximab-treated and untreated PDXs). Access to these data will be granted upon registration to EGA and request to access these studies. Processed expression levels and raw read counts are publicly available in GEO (GSE204805). Supplementary Table 12 lists the results of the targeted sequencing and Supplementary Table 14 reports the GISTIC analysis of shallow sequencing data. Drug responses are presented in Supplementary Table 6. The whole RNAseq dataset deposited in the EGA and GEO repositories comprises 210 samples from primary CRC tumors that were not analyzed in this manuscript, but were processed for overall quality checks and principal component analyses because they were sequenced in the same batches. Source data files are available for Figures 1, 5, 6 and Supplementary Figures 1, 12 and 13. For the remaining figures, all other intermediate analysis data are available from the corresponding authors upon request.

### Code availability

All pipelines used to parse and analyze DNA and RNA sequencing data were developed and run with snakemake (v5.4.0); the code is available at the following repositories: i) https://github.com/vodkatad/snakegatk (targeted sequencing and shallow sequencing alignment and GATK best practices); ii) https://github.com/vodkatad/RNASeq_biod_metadata (RNAseq overall QC and metadata management); iii) https://github.com/vodkatad/biodiversa_DE (RNAseq differential analysis); iv) https://github.com/vodkatad/biobanca (overall comparative analyses).

## ACKNOWLEDGMENTS

We acknowledge Merck KGaA for providing cetuximab. We thank Mauro Papotti, Gianluca Paraluppi, and Serena Perotti for sample acquisition; Barbara Martinoglio for support with real-time PCR; Alessandro Fiori and Massimiliano Frassà for data management; Massenzio Fornasier and Arianna Russo for veterinary assistance; Fabrizio Maina for animal husbandry; Raffaella Albano, Stefania Giove, Lara Fontani, and Laura Palmas for technical assistance; and Daniela Gramaglia and Mauro Paschetta for secretarial assistance. This work was conducted with funding from AIRC, Associazione Italiana per la Ricerca sul Cancro, Investigator Grants 20697 (to A.B.), 22802 (to L.T.), and 23211 (to L.P.); AIRC 5×1000 grant 21091 (to A.B. and L.T.); AIRC/CRUK/FC AECC Accelerator Award 22795 (to L.T.); AIRC MFAG 25040 (to A.P.); European Research Council Consolidator Grant 724748 BEAT (to A.B.); H2020 grant agreement no. 754923 COLOSSUS (to L.T.); H2020 INFRAIA grant agreement no. 731105 EDIReX (to A.B.); Horizon Europe grant agreement no. 101058620 canSERV (to L.T.); and Fondazione Piemontese per la Ricerca sul Cancro-ONLUS, 5×1000 Ministero della Salute 2016 (to L.T.). A.B. and L.T. are members of the EurOPDX Consortium.

---

**Supplementary Figure 1.**
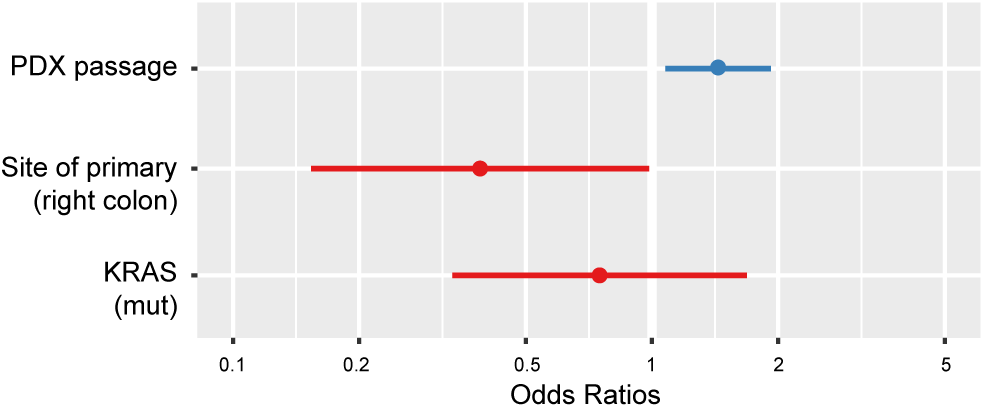
Impact of PDX passage on PDXT validation rates. Odds ratios of a multivariate logistic regression with PDXT validation success status (1, successful, *N* = 64; 0, failed, *N* = 49) as dependent variable. PDX passage at which tumoroids were derived is the independent variable, together with the two significant enrichments shown in Fig. 1F (*KRAS* mutation and tumor right sidedness). When PDX passage is computed in the multivariate analysis, the odds ratio of tumor sidedness maintains statistical significance, whereas the odds ratio of *KRAS* mutations becomes not significant. This suggests that *KRAS* mutant PDXTs were preferentially derived from early-passage xenografts, which were more prone to fail the validation procedure. PDX passage is a continuous variable; site of primary and *KRAS* mutations are binary variables.

**Supplementary Figure 2.**
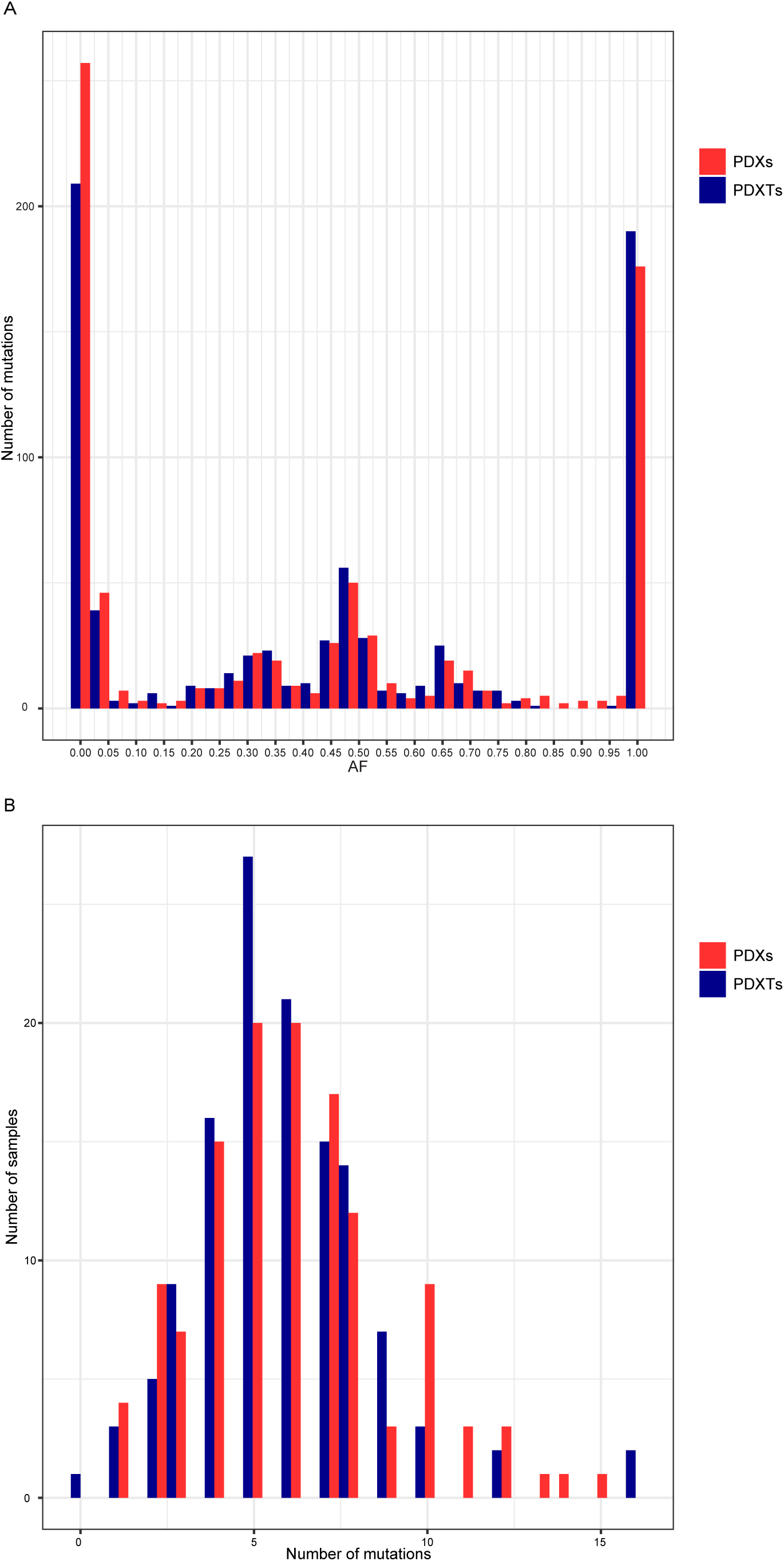
General overview of targeted sequencing results in matched PDXs and PDXTs. **A,** Distribution of mutation allele frequencies in a set of 144 PDXTs and their matched PDXs. **B,** Number of mutations per sample in the same set.

**Supplementary Figure 3.**
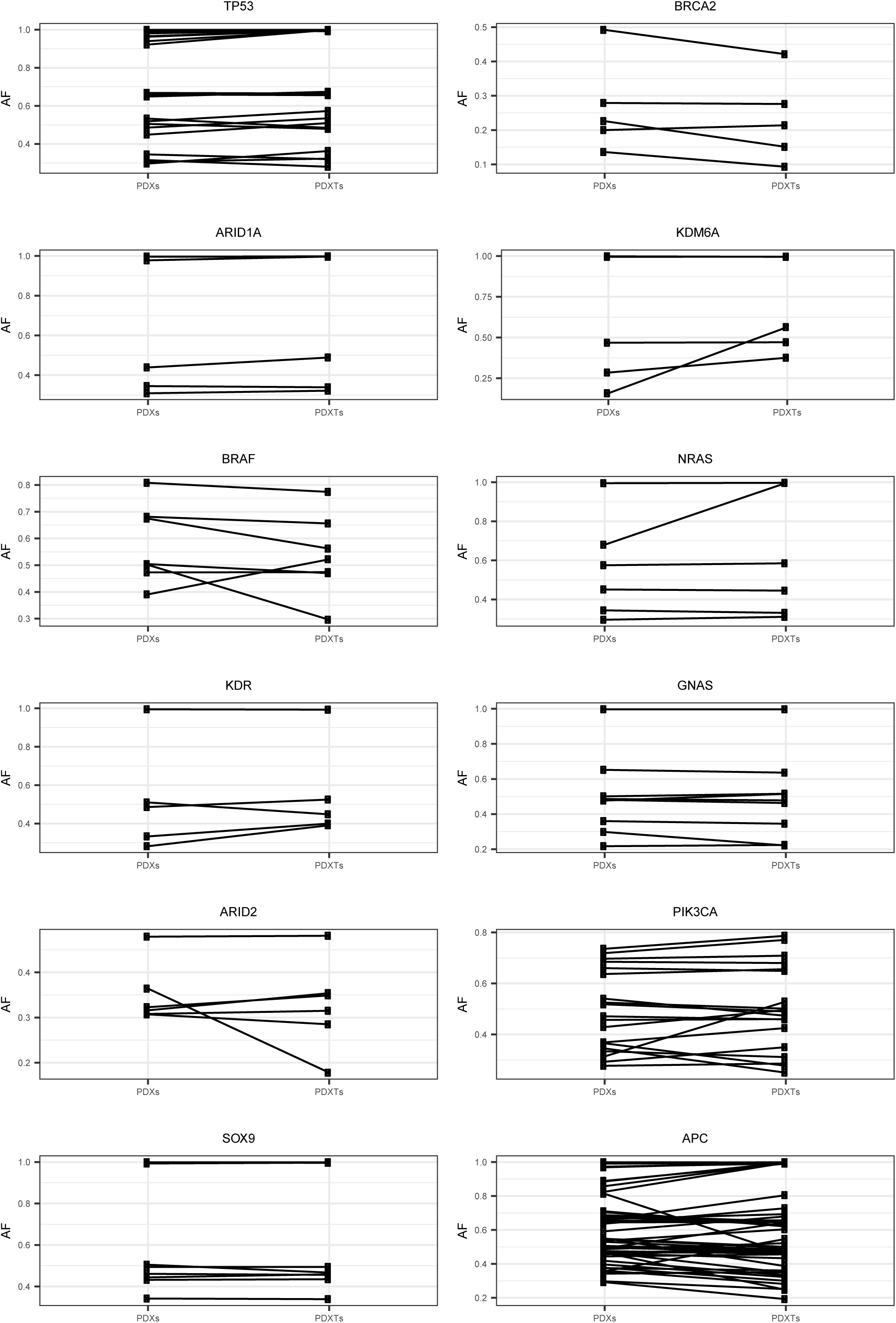
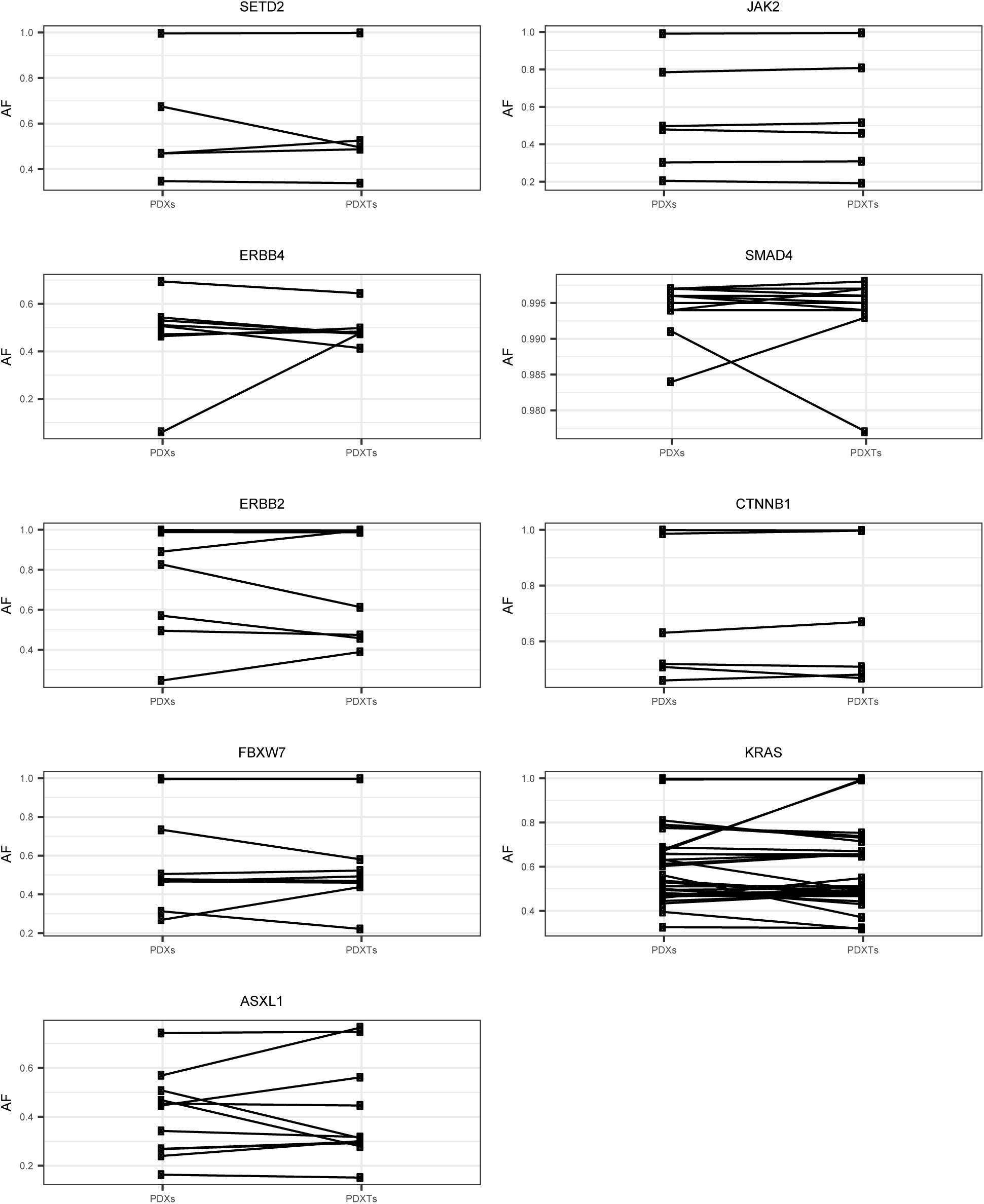
Clonal representation of shared mutations in matched PDXs and PDXTs. Comparison of allele frequencies (threshold, 0.05) for genes with more than five alterations in the collection. *P* = 0.66 by paired *t*-test. AF, frequencies.

**Supplementary Figure 4.**
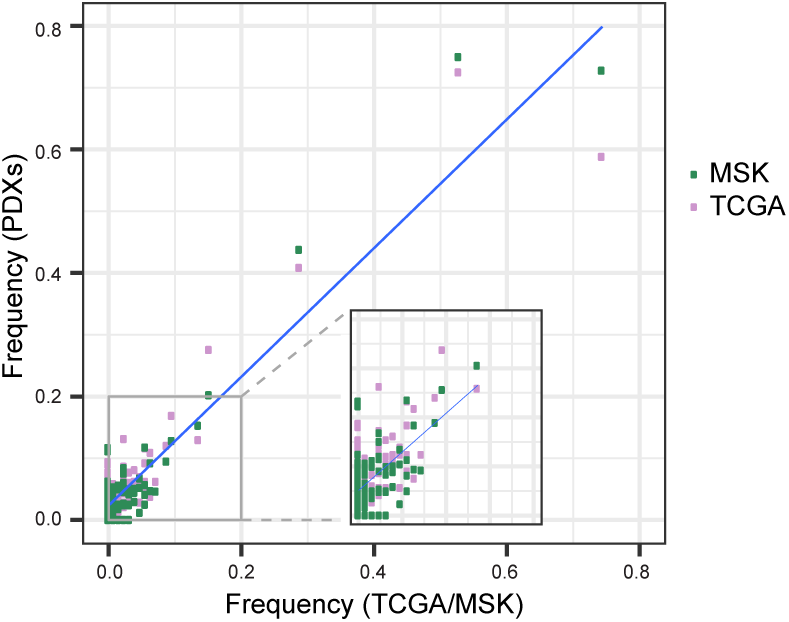
Frequencies of mutant genes in PDXs versus public datasets. Gene-level population frequencies of mutational alterations in PDXs versus those detected in the TCGA dataset or the MSK-IMPACT dataset; the inset shows that the correlation is not driven solely by genes with high mutational frequencies. Pearson coefficient, 0.92 (*P* = 4.24e-49) for TCGA; Pearson coefficient, 0.95 (*P* = 4.57e-62) for MSK-IMPACT.

**Supplementary Figure 5.**
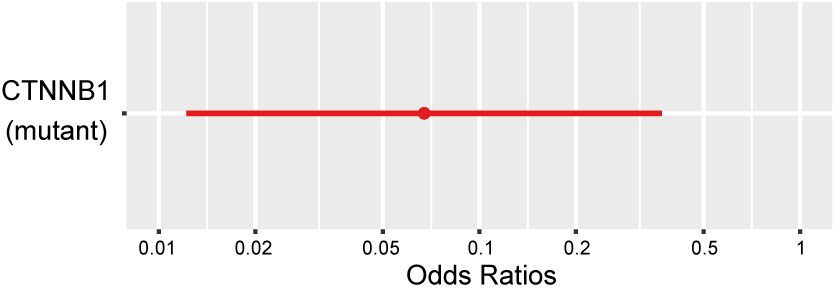
Impact of β-catenin (*CTNNB1*) mutational status on PDXT validation rates. Odds ratios of a multivariate logistic regression with PDXT validation success status (1, successful, *N* = 125; 0, failed, *N* = 7) as dependent variable and *CTNNB1* mutational status as independent variable.

**Supplementary Figure 6.**
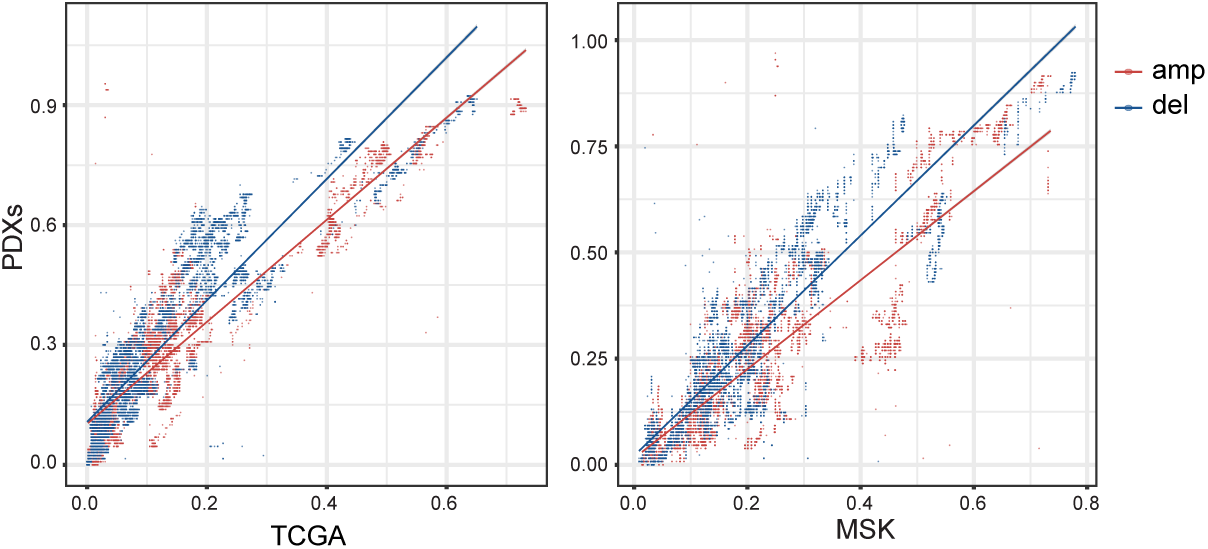
Frequencies of genes with copy number events in PDXs versus public datasets. Gene-level population frequencies of gain or loss events, as identified by GISTIC, in PDXs versus those detected in the TCGA or the MSK-IMPACT datasets. Pearson coefficient, 0.92 (gains) and 0.90 (losses) for TCGA; 0.84 (gains) and 0.89 (losses) for MSK-IMPACT (*P* < 1e-230 for all comparisons). Amp, amplification; del, deletion.

**Supplementary Figure 7.**
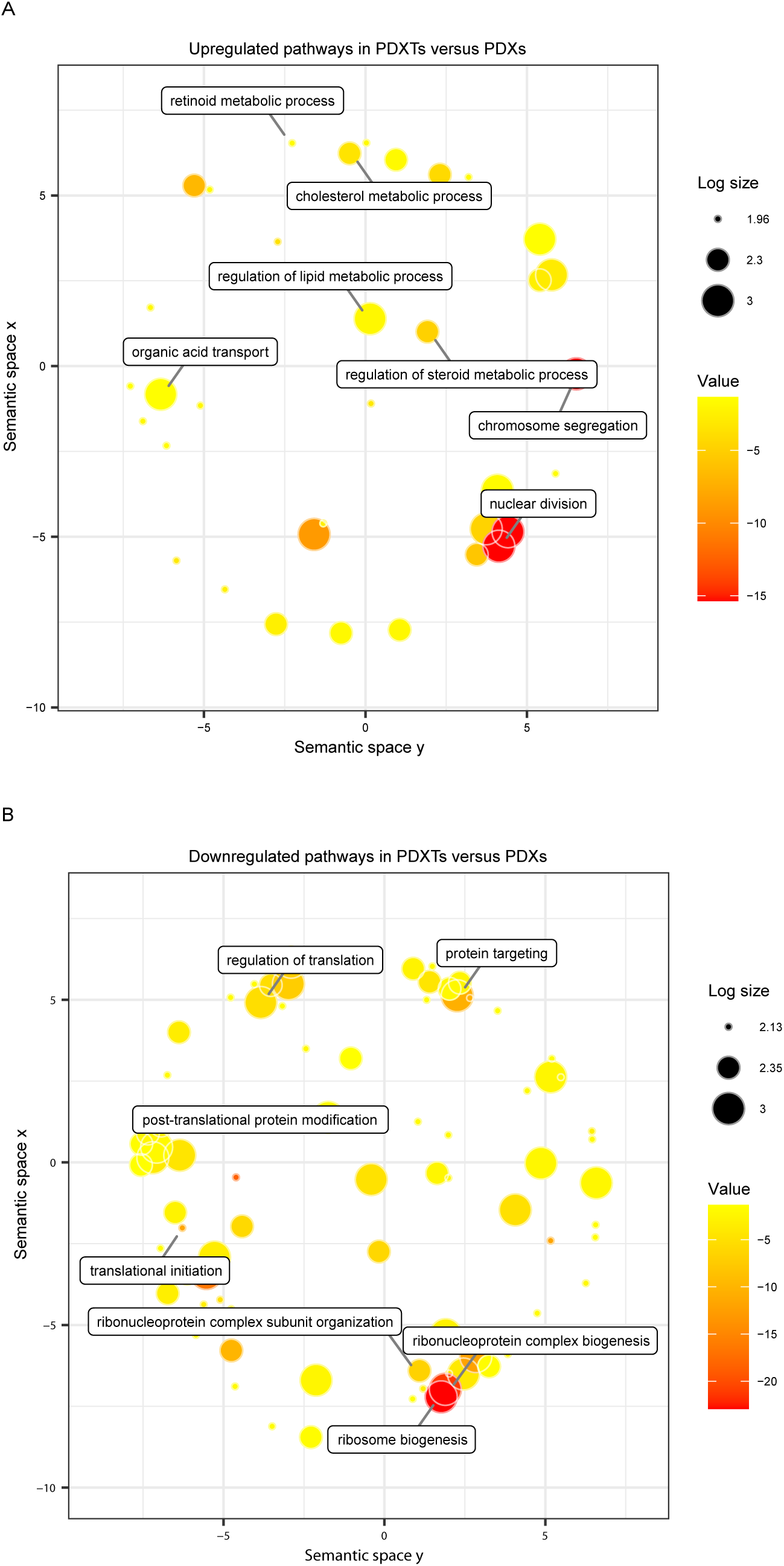
Differentially modulated pathways in PDXTs versus matched PDXs. GO (Biological Processes) enrichment analysis for pathway signatures that are differentially expressed in PDXTs and matched PDXs. GO terms were visualized with REVIGO^71^ The size of the circles reflects the number of genes annotated with one GO term (log10 scale); the color denotes significance of the enrichment (log10 scale). The Log size legend shows the median, the third quartile and the maximum values. Circle positioning in the 2D space signifies the semantic similarity between terms. When GO definitions are very similar, only one term is selected by REVIGO.

**Supplementary Figure 8.**
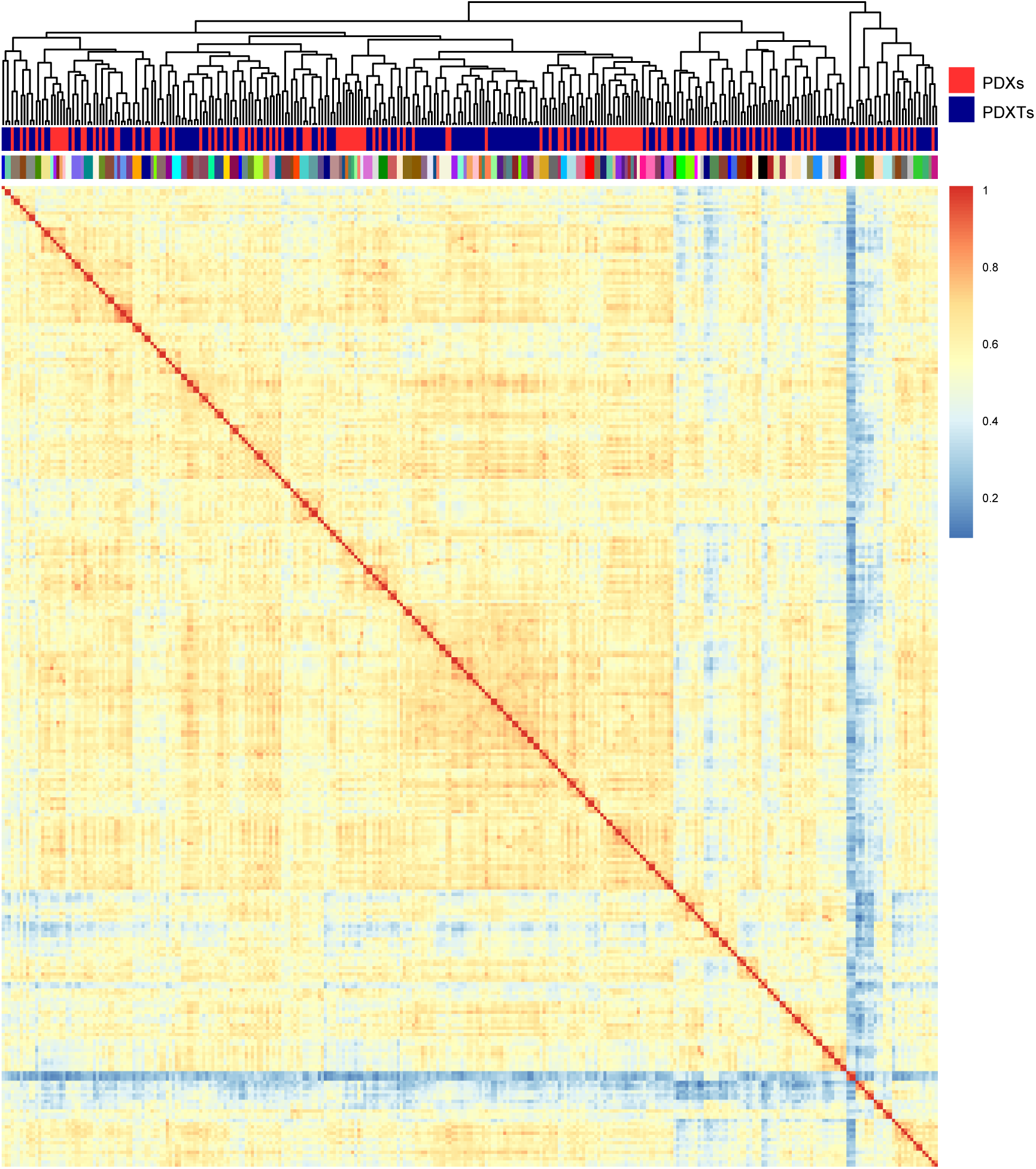
Global comparison of transcriptional profiles in matched PDXs and PDXTs. Hierarchical clustering of matched PDXs and PDXTs based on the Pearson correlation between their expression profiles. The samples analyzed (*N* = 308) include some biological replicates (in the case of PDXs, the same tumor engrafted in different mice; in the case of PDXTs, the same tumor propagated in different plates and/or recovered after different freeze-thaw cycles). Each tumor of origin is identified by a specific color in the top annotation.

**Supplementary Figure 9.**
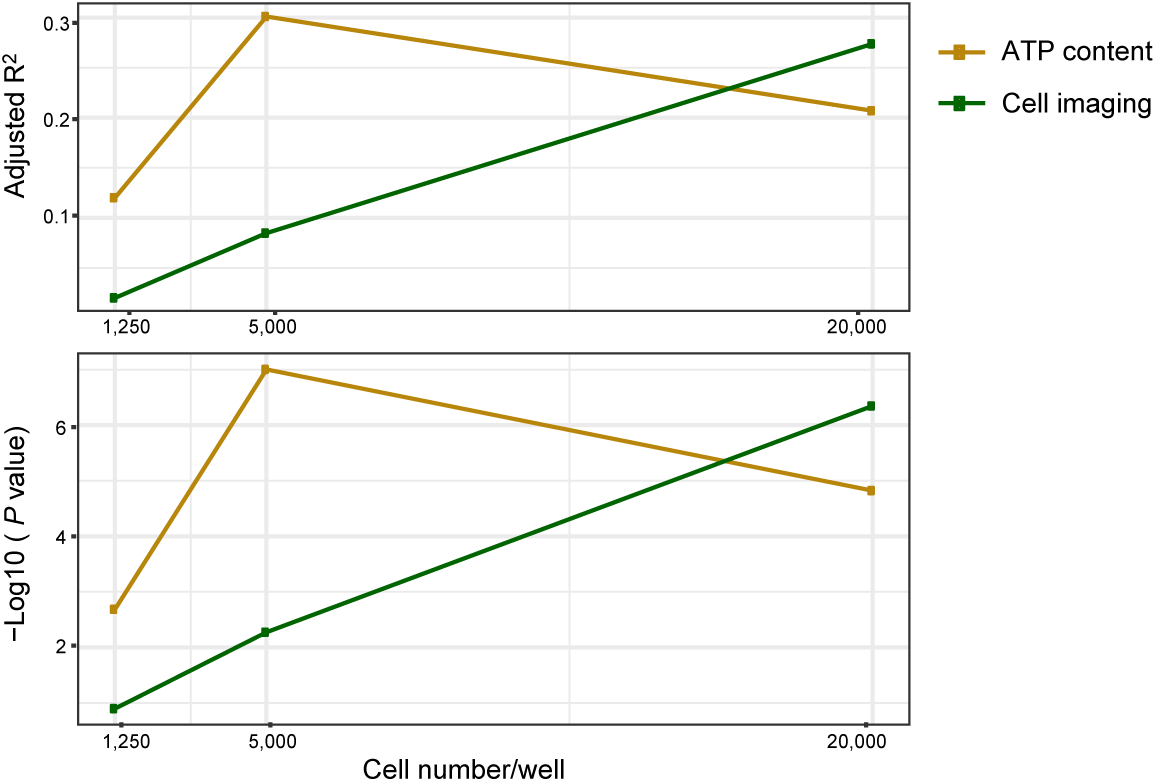
Performance of PDXT cellular densities in predicting cetuximab response in matched PDXs. Adjusted R-squares and -Log10(pvalue) of linear regression models with PDX response as dependent variable and different plating densities (cell number/well) and readouts (ATP content and cell imaging) as independent variables. Adjusted R-squares are needed to perform a fair comparison between the different plating conditions, since the number of available data points is not the same for all conditions (69 for 1250 cells/well; 79 for 5,000 cells/well and 20,000 cells/well).

**Supplementary Figure 10.**
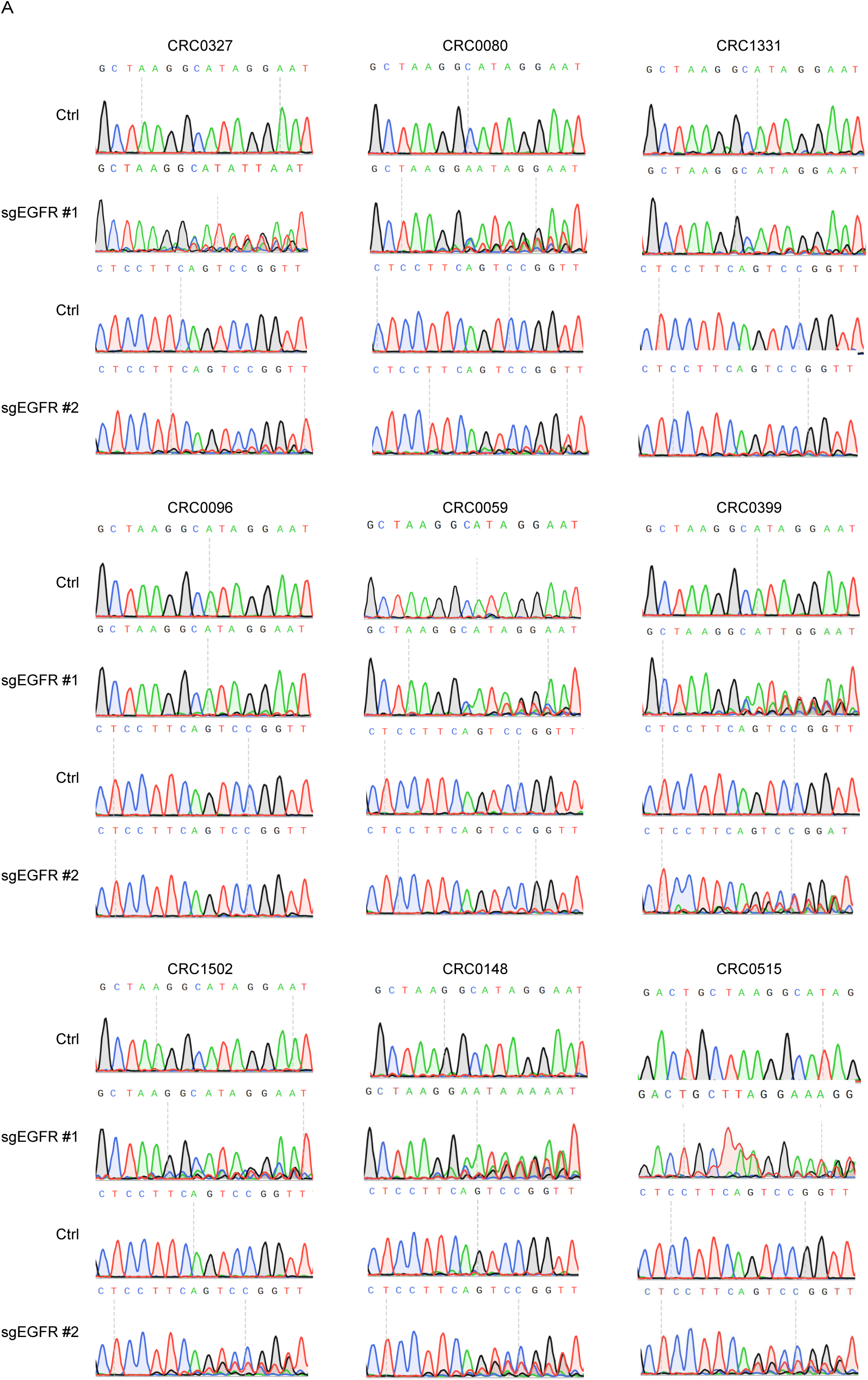
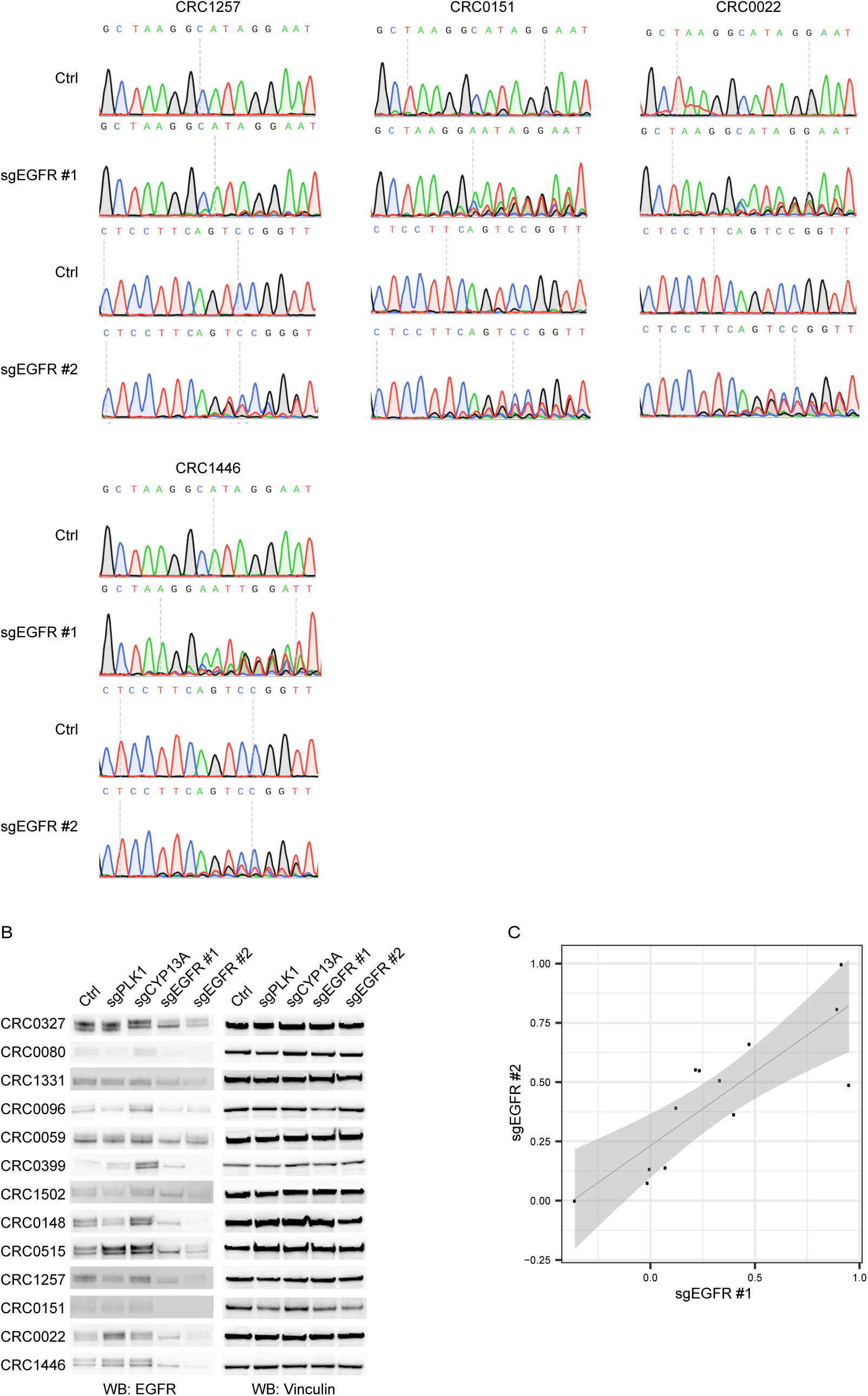
CRISPR-Cas9-mediated inactivation of EGFR in PDXTs. Cas9-expressing PDXTs (control) were transduced with the following guides: i) two different sgRNAs targeting *EGFR* (sgEGFR #1 and sgEGFR #2); ii) a sgRNA targeting the essential gene *PLK1* (sgPLK1); iii) a sgRNA targeting the non-essential gene *CYP13A* (sgCYP13A). Four days after transduction, DNA and proteins were extracted in parallel to evaluate *EGFR* gene disruption and expression, respectively. **A,** Sanger sequencing chromatograms of CRISPR/Cas9 genome-edited cell populations, showing the expected genomic scar in *EGFR* exon 3 sequence. Wild-type exon 3 sequence of the corresponding control models was used as control. Each image is representative of two independent experiments. **B,** Western blot analysis of EGFR protein expression. Vinculin was used as loading control. Each image is representative of two independent experiments. **C,** Linear regression of EGFR KO scores using sgEGFR #1 or sgEGFR #2. Results are the average of two independent experiments, each performed in biological triplicates (with the exception of CRC0148, which was tested in three independent experiments). Pearson, 0.84 (*P* = 3.3e-4). Ctrl, control.

**Supplementary Figure 11.**
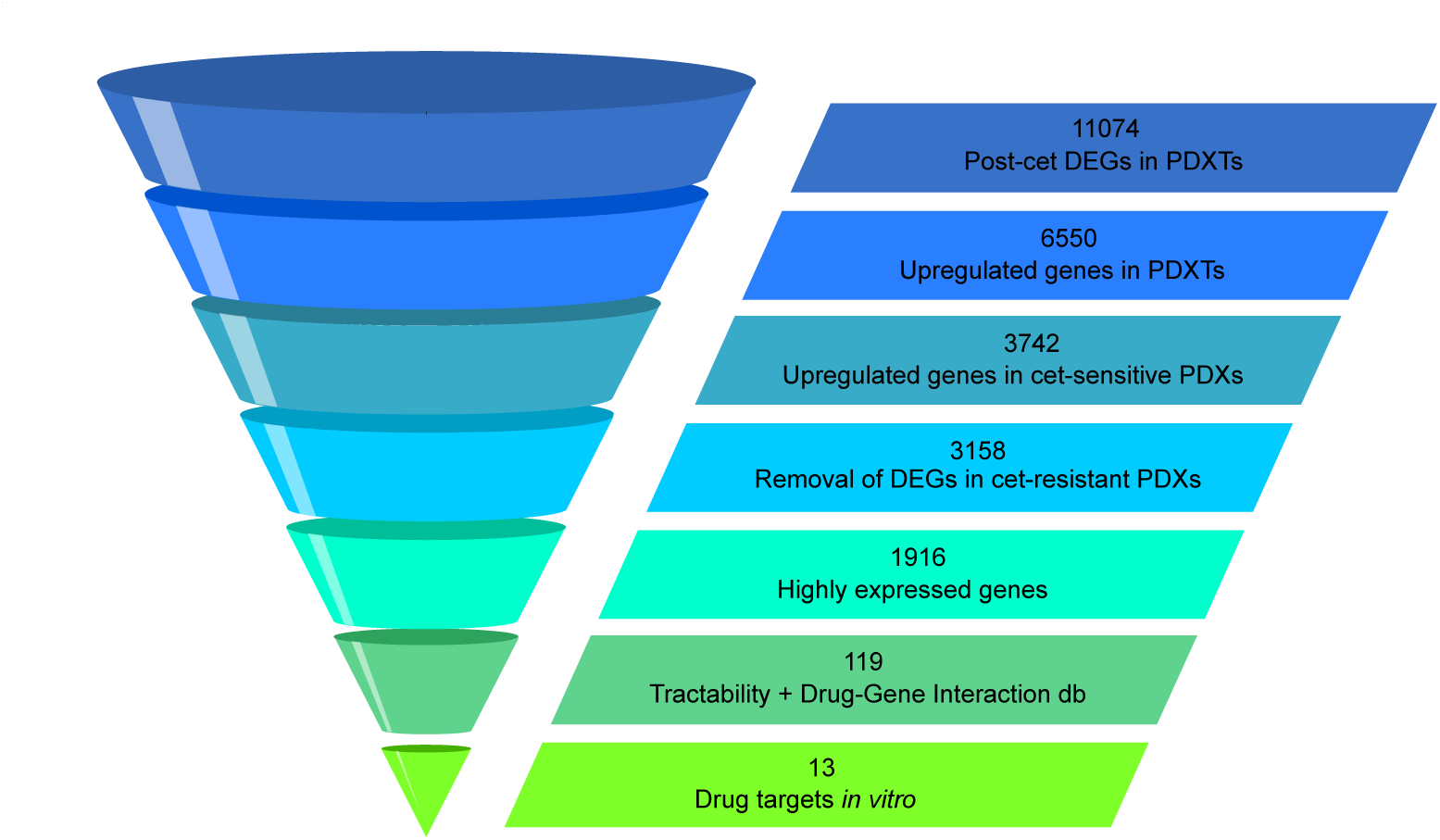
Stepwise prioritization of cetuximab co-extinction targets. To extract biologically meaningful targets adaptively upregulated by EGFR inhibition, the list of genes modulated by cetuximab in PDXTs was restricted by applying sequential filters, including: i) the identification of genes concordantly upregulated by antibody treatment in cetuximab-sensitive PDXs; ii) the removal of genes showing expression changes after antibody treatment in cetuximab-resistant PDXs; and iii) the selection of highly expressed genes. The resulting genes were further shortlisted through an educated assessment of target tractability and a search of druggable targets in the Drug-Gene Interaction Database (www.dgidb.org). Cet, cetuximab; db, database; DEGs, differentially expressed genes.

**Supplementary Figure 12.**
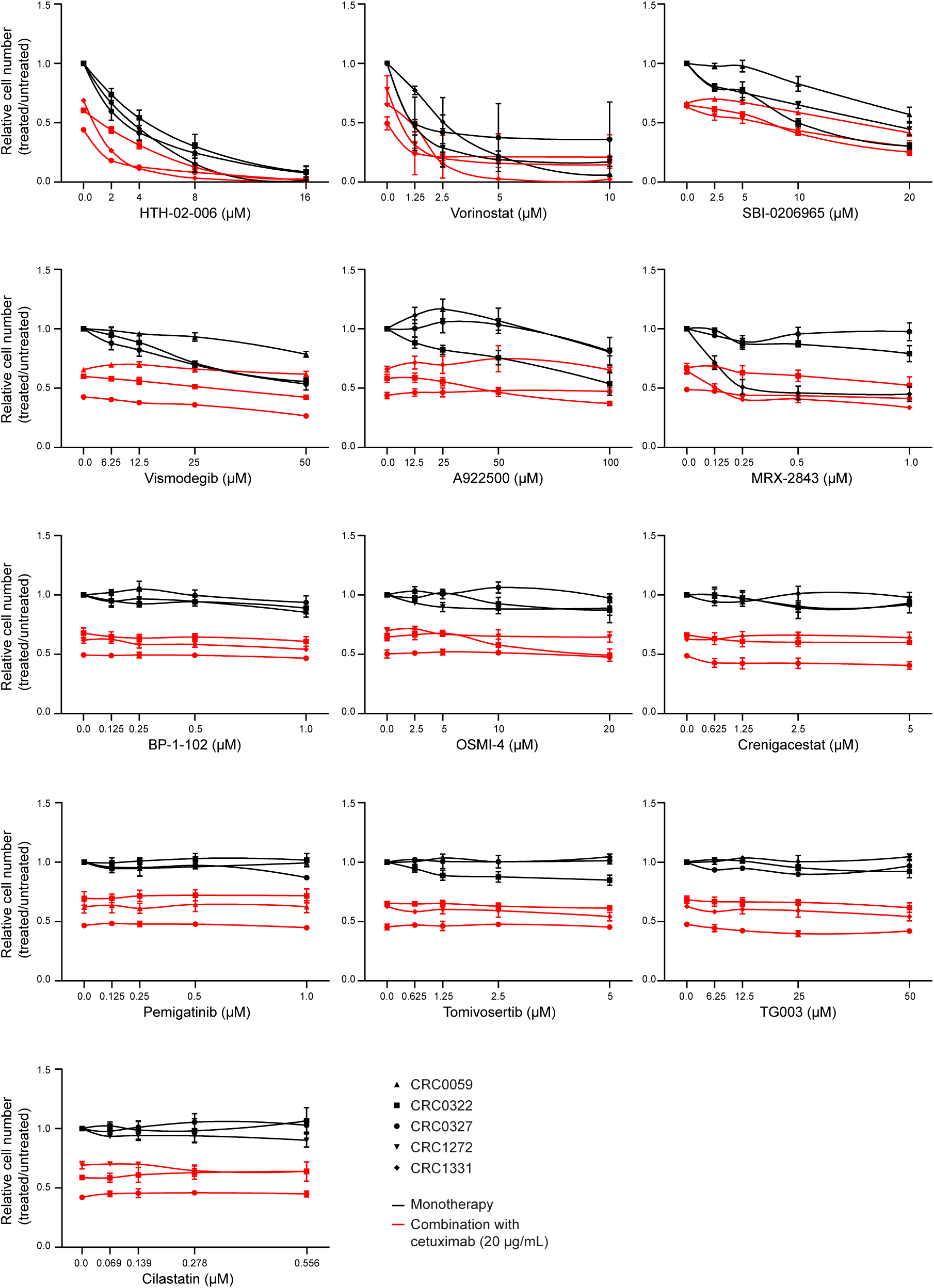
Short-term activity of selected compounds, alone or in combination with cetuximab, in PDXTs. Relative cell viability (ATP content) in PDXT models treated with the indicated modalities for 48 hours. Results represent the means ± SD of three independent experiments performed in biological triplicates (*N* = 9). Results of statistical analysis (one-way ANOVA) are reported in Extended Data Table 9.

**Supplementary Figure 13.**
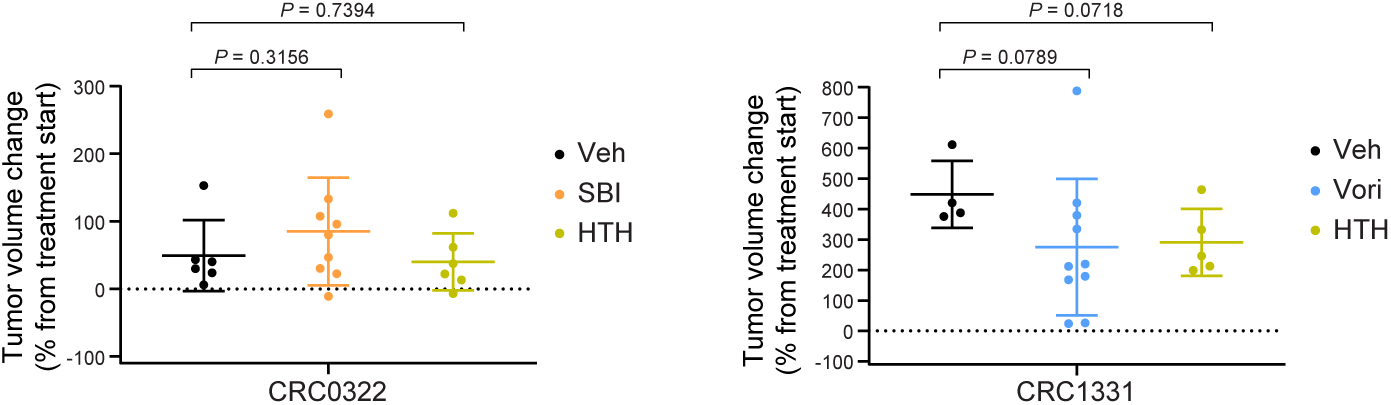
Single-agent activity of selected compounds in PDXs. Tumor volume changes in PDXs from mice treated for one week (CRC0322) or three weeks (CRC1331) with vehicle (until tumors reached a maximum tumor diameter ≥ 10 mm), HTH-02-006 alone (10 mg/kg, intraperitoneal injection twice a day), SBI-0206965 (20 mg/kg, intraperitoneal injection three times a week), or vorinostat (50 mg/kg, intraperitoneal injection three times a week). In the case of CRC0322, the experiment was terminated at one week because some mice had to be euthanized upon reaching of the humane endpoint. Dots represent volume changes of PDXs from individual mice, and plots show the means ± SD for each treatment arm. *N* = 4 to 10 animals per each treatment arm. Statistical analysis by two-tailed unpaired *t* test with Welch’s correction. HTH, HTH-02-006; SBI, SBI-0206965; Veh, vehicle; Vori, Vorinostat.

**Supplementary Figure 14.**
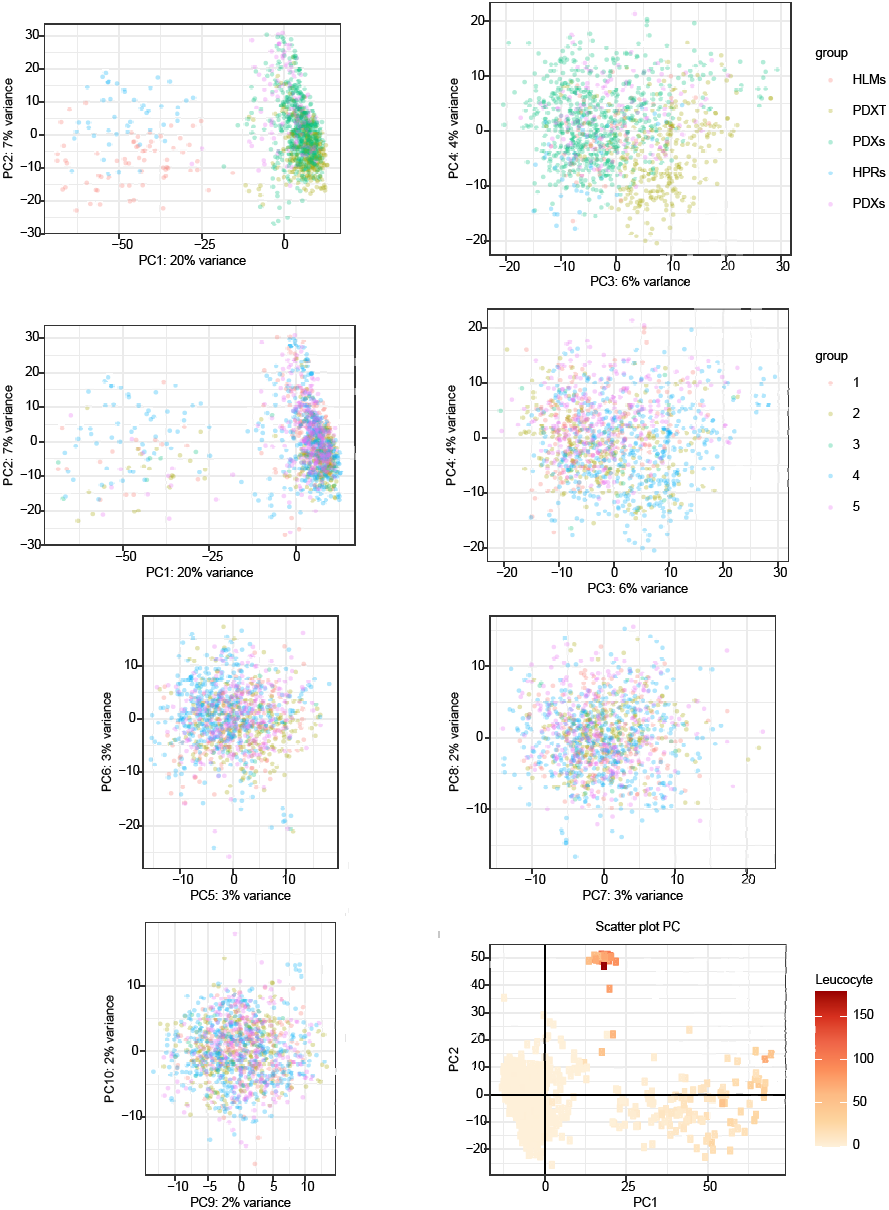
Overall representation of the RNAseq dataset via Principal Component Analysis. A-C, Main variations in expression values, represented with principal components (PC), with respect to (A) sample origin (PC1, PC2, PC3 and PC4) and (B) batches (PC1 to PC10). C, PC1 and PC2 with single dots shaded according to the leucocyte expression signature. HLMs, human liver metastases; HPRs, human primary tumors.

